# A Genome-Wide Comprehensive Analysis of Nucleosome Positioning in Yeast

**DOI:** 10.1101/2023.06.02.543396

**Authors:** Leo Zeitler, Kevin André, Adriana Alberti, Cyril Denby Wilkes, Julie Soutourina, Arach Goldar

## Abstract

In eukaryotic cells, the one-dimensional DNA molecules need to be tightly packaged into the spatially constraining nucleus. Folding is achieved on its lowest level by wrapping the DNA around nucleosomes. Their positioning regulates other nuclear processes, such as transcription and DNA repair. Despite strong efforts to study nucleosome phasing using Next Generation Sequencing (NGS) data, the mechanism of their collective arrangement along the gene body remains poorly understood. Here, we assess the nucleosome profiles of protein-coding genes in *Saccharomyces cerevisiae* using functional Principal Component Analysis. By decomposing the NGS signals into their main descriptive functions, we compared wild type and chromatin remodeler-deficient strains, keeping position-specific details preserved. A correlation analysis with other genomic properties, such as gene size and length of the upstream Nucleosome Depleted Region (NDR), identified key factors that influence nucleosome phasing. We reveal that the RSC chromatin remodeler—which is responsible for NDR maintenance—is indispensable for decoupling nucleosome arrangement within the gene from phasing outside, which interfere in *rsc8*-depleted conditions. Moreover, positioning in *chd1*Δ strains displayed a clear correlation with RNA polymerase II presence, whereas wild type cells did not indicate a noticeable interdependence. We propose that RSC is pivotal for global nucleosome organisation, whilst Chd1 plays a key role for maintaining local arrangement.

## Introduction

The eukaryotic DNA must be tightly wrapped into the spatially constraining nucleus. This is achieved in the form of chromatin, a DNA-protein complex within which the 1-dimensional DNA is condensed around histone octamers and folded to a 3-dimensional structure. To be more precise, these histone complexes are positively-charged multiprotein structures around which the DNA molecule is locally coiled, forming a linear organisation resembling the stringing together of beads. This is why the primary structure of chromatin is commonly represented by a so-called *beads-on-a-string* model. In yeast, a nucleosome refers to *≈*147 base pairs (bp) of DNA that is wrapped around four histone units. Nucleosomes are closely spaced, with an averaged center-to-center distance of 165 bp, leaving roughly 15 bp of linker DNA between two adjacent histone complexes. There is a consensus that phasing is highly regular within coding regions, which is shortly interrupted by Nucleosome Depleted Regions (NDRs) between two neighbouring genes. This observation gave rise to the barrier model, which proposes that promoter-dependent properties (e.g. bound proteins or sequence composition) pose a limit for nucleosome assembly, and arrangement occurs with respect to this barrier [1, 2]. However, it is widely accepted that various factors establish and influence the genome-wide positional nucleosome landscape, including sequence composition, transcription, and chromatin remodelers [3–6]. Since the DNA molecule must bend to wrap around the histone octamer, the local nucleotide sequence naturally affects positioning. Generally speaking, GC-rich sequences are more flexible than AT-rich ones, and they are favorable to support the presence of a nucleosome [7, 8]. However, sequence-related properties might be dependent on specific motifs.

The condensed packaging also functions as regulator for various DNA-protein interactions. Most of these processes rely on chromatin remodeler complexes, which can—by consuming energy obtained from ATP hydrolysis—move, add, or evict the histone complexes to provide or inhibit direct access to the DNA sequence [9]. In yeast, chromatin organisation is maintained by four protein families, SWI/SNF, INO80, ISW and CHD. The RSC remodeler complex of the SWI/SNF family is the only essential chromatin remodeler in *Saccharomyces cerevisiae*, and it is recruited to promoter regions where it is responsible for the maintenance of NDRs [10–12]. The complex is also reportedly influencing nucleosome organisation in coding regions as well as supporting RNA Polymerase II (Pol II) elongation [13]. It is presumed to restore chromatin organisation after transcription [14]. However, RSC does not exhibit an impact on regular nucleosome spacing within the gene [14, 15]. Chd1—the only member of the CHD remodeler family in yeast—is associated with various transcription-regulating functions, including initiation, elongation, and termination [16]. It has been suggested that Chd1 stabilises perturbed nucleosomes during gene expression [17]. Isw1 and Chd1 are supposed to antagonise for nucleosome spacing within the gene, with Isw1 dominating genes with larger spacing, whereas Chd1 seems to control shorter spacing [12, 18]. Indeed, it has been reported that deletion of Chd1 and Isw1 only disrupt inter-nucleosome distances and leave the +1 position unaffected [19]. Isw2 is similarly associated with regular spacing [20], and it is particularly affecting nucleosomes close to the NDR, which is presumed to regulate transcription [21]. However, the underlying mechanism for chromatin remodelers is still under debate, and a scientific consensus is missing [22–25].

Several studies showed an interdependence between nucleosome distribution and gene expression by using MNase-seq data, a Next Generation Sequencing (NGS) technique that allows the measurement of nucleosome profiles by using MNase digestion of purified chromatin [26, 27]. It has been suggested that high gene expression correlates with low nucleosome regularity, which is characterised by weak phasing and extreme spacing (both short and long) [18, 28]. Furthermore, promoter regions tend to be more open when transcription levels are high [29]. Indeed, the depletion of Pol II exhibited increased array regularity [30]. This phenomenon seems to be conserved across species, as indicated by studies using *Drosophila* [31] and mouse cell lines [32]. The outcomes indicate that gene expression can be partially explained by nucleosome positioning over the gene body. Nonetheless, the autocorrelation of MNase-seq profiles along genes revealed that nucleosomal organisation accounts for only *≈*25% of the observed transcriptional variability, even though genes with similar regularity tend to have the same level of gene expression [33]. Surprisingly, many strains deficient for chromatin remodelers seem to show only a marginal effect on transcription [18, 19]. The only exception is *rsc8*-depleted cells, which exhibit a global decrease in gene expression [12]. A clear picture between nucleosome phasing and Pol II presence is still lacking.

Different approaches have been used to categorise collective nucleosome arrangement within transcribed regions. However, many of them rely on Pearson [34] or autocorrelation measurements [33], both of which only allow to quantify the average behaviour over all considered nucleosomes. Another analysis that takes into account multiple nucleosomes upstream and downstream of the NDR was presented by [14]. However, the study focused on changes with respect to the NDR, and many phenomenological descriptions are based on the application of different analysis techniques. To our knowledge, a single mathematical framework assessing gene-wide nucleosome phasing has not been proposed, and a direct comparison of the effects in different remodeler-deficient strains is difficult.

In this work, we present a genome-wide analysis of nucleosome positioning along the gene. To be precise, we assess the location-specific arrangement of 6-7 nucleosomes into the gene body. By doing so, we address two points that have previously largely fallen short. Firstly, we evaluate long-range influences of positioning within transcribed regions that exceed neighbouring nucleosomes in a data-driven manner; and secondly, we analyse local effects necessary for coordinating nucleosome arrangement as *beads on a string*. Our approach therefore refines and complements other studies that focused on a few individual nucleosomes close to the NDR or Transcription Starting Site (TSS); or which assessed only the average correlation of the entire array. We combined a conventional linear Pearson correlation measurement with functional Principal Component Analysis (fPCA) to investigate the functional composition of nucleosome profiles on a genomic scale. FPCA is commonly used in time series and signal processing, and it has been used in biology for analysing crop yield [35], identifying child growth patterns [36], as well as studying genetic variation and the allelic spectrum [37]. However, it has never been applied to the spatial interdependence of nucleosome phasing to our knowledge. By using MNase-seq data from yeast strains deficient for different chromatin remodelers [12, 18], we reveal that Rsc8 strongly limits coordinated nucleosome arrangement to the transcribed region. It might be therefore responsible for gene-specific phasing. Our methods allow quantifying the effect of several remodelers on nucleosome positioning. We identified 5 combinations of gene deletions or protein depletions which have a notable impact on phasing compared to Wild Type (WT) conditions. Measuring correlation with other nuclear processes disclosed that none of the commonly assumed factors can easily explain long-reaching nucleosome arrangement in WT strains within the gene body. However, gene deletions—in particularly mutants that contained *chd1*Δ—caused a strong correlation with Pol II presence. Our results indicate a new mechanistic understanding of chromatin remodelers, where Rsc8 is responsible for long-range coordination and Chd1 for local positioning of nucleosomes.

## Results

### Nucleosome Profiles Can Be Well Distinguished Based On Their Coordinated Positioning in WT

In order to compare nucleosome profiles over the gene body in WT conditions, we measured the Pearson cross-correlation of the MNase-seq data produced by [12, 18] for all protein-coding regions [38]. This follows previous analyses using similar measurements [33, 34]. For both replicates, we considered 1000 bp after and 200 bp before the +1 position (= 1200 bp, approximately the average size of a gene in *Saccharomyces cerevisiae*), containing 6-7 nucleosome dyads. Pearson coefficients were grouped into two distinct partitions using *k*-mean clustering. By comparing the Jensen-Shannon (JS) distance of the Pearson clusters with 500 random groups, we proved their significance (outside the 95% prediction interval (PI) of a gamma distribution estimated over the random partitions; SFig A.1). This shows that nucleosomal arrays can be significantly separated into two groups by their linear correlation.

It is worthwhile for the following analysis to explain what the two Pearson clusters represent. As nucleosomes are commonly well positioned in budding yeast, the MNase-seq data resembles a wave-like function with one peak approximately every 200 bp. The Pearson correlation expresses therefore average similarity of *coordinated nucleosome positioning* between two profiles. The amplitude is not taken into account. Each group contains all those distributions that are most similar to each other. *Coordinated positioning* refers to the configuration of the entire nucleosome array; thus, phasing of a single nucleosome needs to be considered with respect to all others. Due to the wave-like form of the signal, differences in similarity come either from shifts in exact positioning (i.e. well-defined peaks) or from a general change in the signal amplitude (i.e. increasing or decreasing MNase-seq magnitude over the profile). The clusters are separated based on these trends (Fig 1(A)). It should be emphasised that the link between linear correlation and coordinated nucleosome phasing can only be established due to the repetitive wave-like function where each peak represents a single nucleosome position. For other distributions—where this distinction is not clear—Pearson correlation would require a different interpretation.

**Figure 1:**
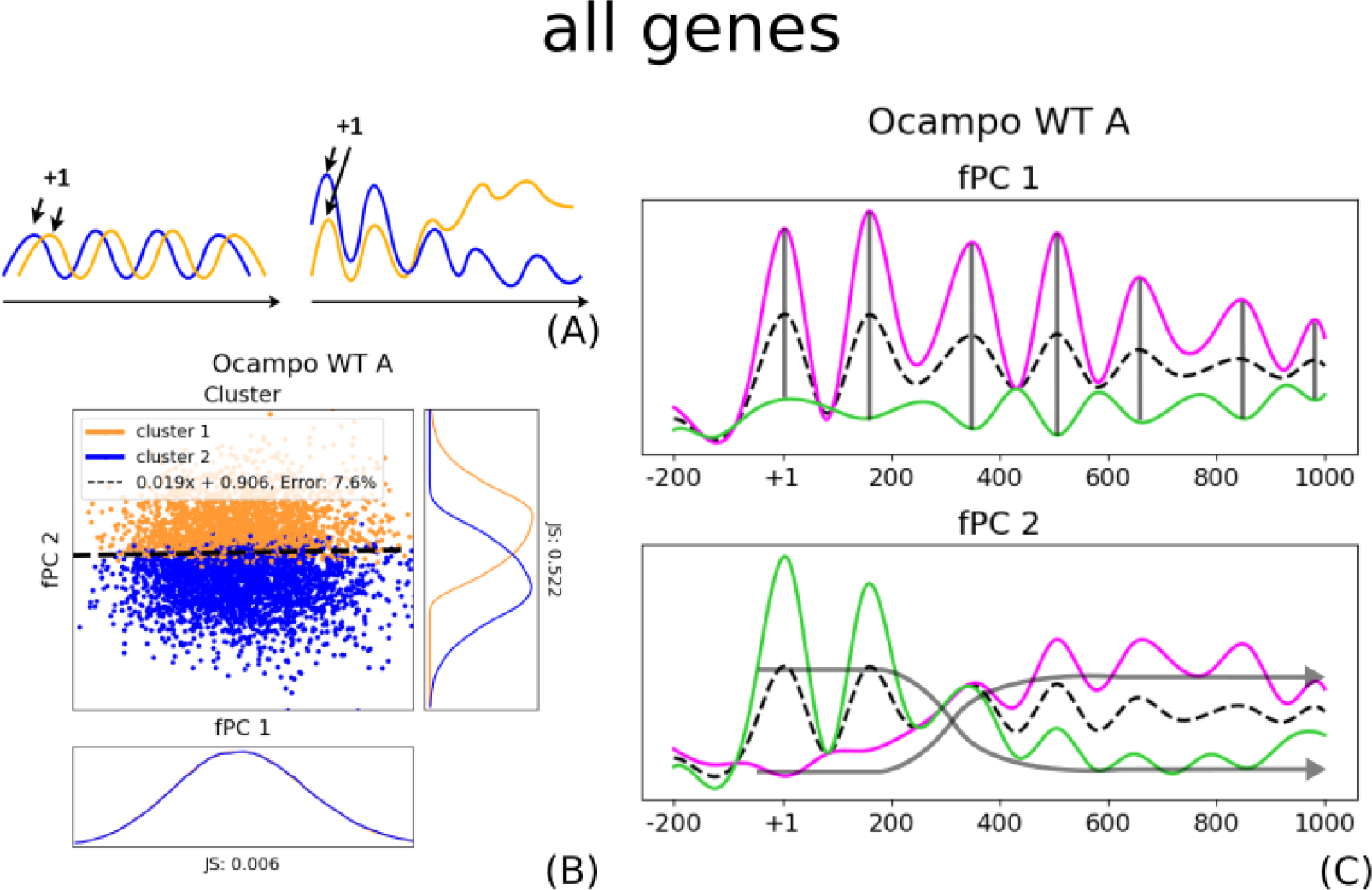
FPCA and Pearson clusters considering all protein-coding genes. (A) Due to the well-positioned nucleosomes and the wave-like structure of MNase-seq data, we presume that the Pearson correlation measures coordinated nucleosome positioning along the gene. If two profiles (orange and blue) are in two different clusters, this could indicate either a shift in the exact nucleosome positions (left); or a general trend in the MNase-seq signal amplitude, i.e. either increasing or decreasing (right). (B) Pearson clusters considering all genes are linearly separable with respect to their fPC scores. The JS distance between the cluster distributions is much larger for fPC 2 than for fPC 1. Orange and blue indicate each one group, the dashed line symbolises the best linear separation using a SVM. The x-axis represents the score of the first fPC *ζ*_1_, the y-axis gives the score for the second fPC *ζ*_2_. Both axes are scaled to the same size. (C) When analysing the effect of the major fPCs, they describe predominantly position-dependent amplitude (transparent black lines, fPC 1) and collective nucleosome phasing (transparent black arrows, fPC 2). The mean is given as a dashed black line, a positive contribution—i.e. adding the fPC to the mean—is displayed in magenta, and a negative contribution—i.e. subtracting from the mean—is shown in green. The second fPC in WT indicate an increasing or decreasing signal magnitude as a function of distance from the TSS, suggesting stronger or weaker presence (corresponding to Figure 1(A, right)).

Importantly, the Pearson coefficient measures only the average linear pairwise correlation over the entire profile, rather than taking position-dependent particularities into account. We assume that the MNase-seq signal can be described as a continuous function, and it can be approximated by a mixture of a finite number of simpler continuous functions. In this study, we use 20 B-splines to represent the MNase-seq data along each gene, which is similar to conventional smoothing techniques. This permitted the application of fPCA to determine the two best-describing functional Principal Components (fPCs) that explain each nucleosome profile. Intuitively, an fPC is a combination of the 20 B-splines together with a score or weight *ζ* indicating their deviance from the mean profile to describe a particular distribution (further explained in Methods). The two functions extend over the entire gene (or rather 1200 bp) and therefore consider location-dependent differences.

Astonishingly, the two clusters—which were independently obtained by classical hierarchical *k*-mean clustering of Pearson coefficients—are neatly separated by the second fPC, whilst they are seemingly independent of the first (Fig 1(B)). This is slightly less clear for the *B* replicate, although still distinct (SFig A.2(B)). As the groups were determined using a linear correlation measurement, we intuited that the second fPC describes coordinated nucleosome phasing along the gene body. We confirmed this by analysing the effect of the second fPC on the function shape. We found that the first fPC largely represents the amplitude at a given position, as it does not influence the location of the peak (Fig 1(C)). To assess the effect on the *beads-on-a-string* organisation within the transcribed region, we focus in the following particularly on changes to the phasing, and therefore, the second fPC. Overall, the analysis shows that position-dependent amplitude and coordinated arrangement are the best two independent functional descriptors that account for most variance between coding regions in MNase-seq data.

### Functional Principal Component Analysis Reveals Size-Dependent Rsc8-Mediated Phasing of Nucleosome Positions

Since the smallest genes are *≈*300 bp long, the 1000 bp window after the +1 position can contain much more than the actual length of the coding region. In order to analyse how nucleosome phasing is affected by the gene size, we repeated the fPCA considering exclusively small (*≤* 1000 bp) or large genes (*>* 1000 bp). If coordinated positioning is substantially affected by the length of the transcribed region, the major two fPCs—which are now estimated dependent on the gene size—should exhibit a changed behaviour with respect to the linear separability. We can confirm that the linear separation is preserved for large genes, although the boundary becomes slightly sloped (Fig 2(A), replicate B SFig A.2(D)). The fPCs determined when considering only large genes are almost identical to the all-gene fPCs (SFig A.3). We therefore presume that the MNase-seq data for large genes can be still best represented using location-dependent amplitude and coordinated nucleosome organisation.

**Figure 2:**
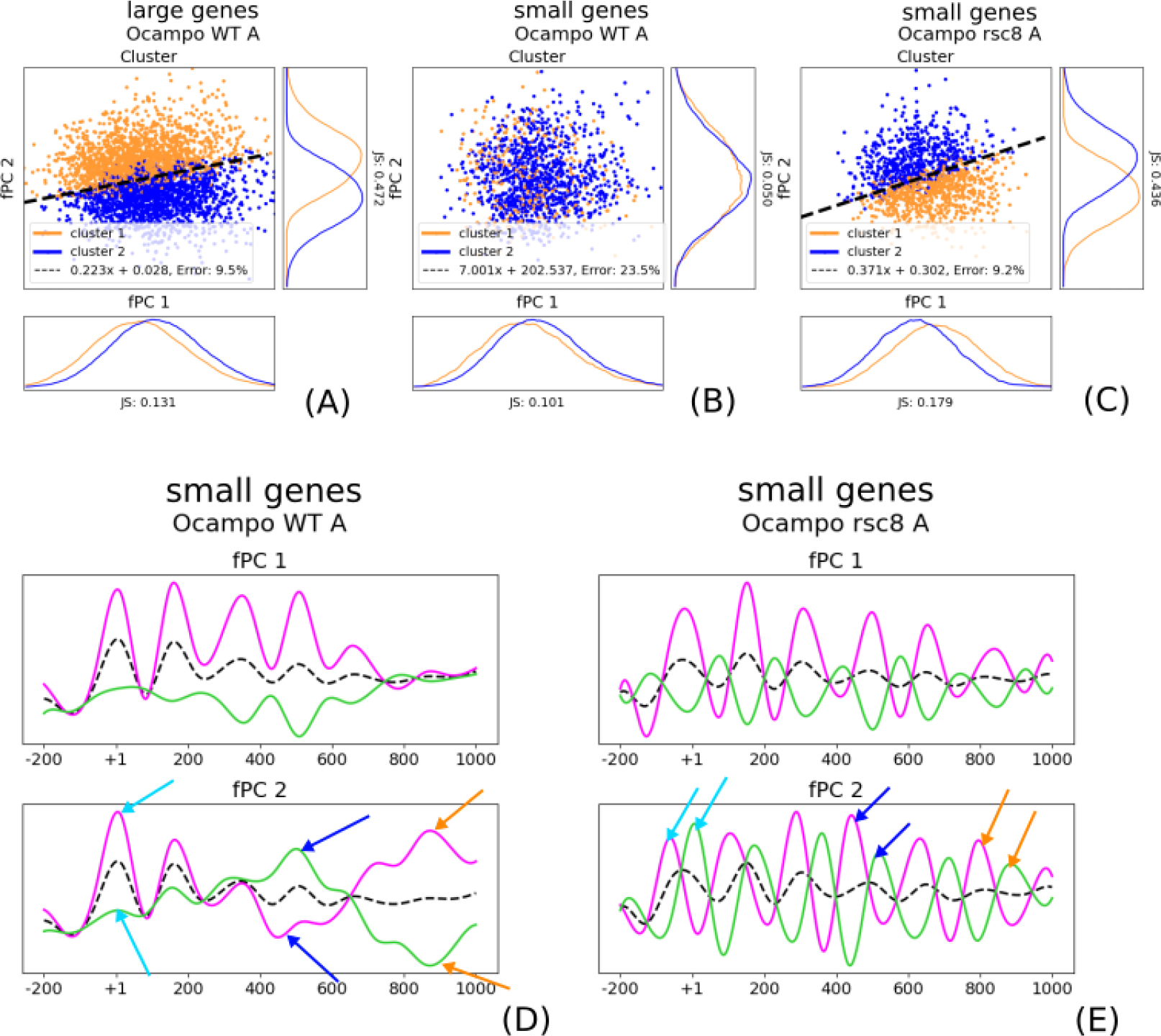
Nucleosome phasing is strictly limited to the gene body, which is maintained by Rsc8. (A) The two Pearson clusters (orange and blue) are still clearly linearly separable (dashed line) when considering exclusively large genes (*>* 1000 bp). (B) However, this boundary is completely lost when performing the fPCA using only small genes (*≤* 1000). Whilst both clusters are present, most short coding regions are in the same Pearson group (blue). (C) Surprisingly, the clusters (blue and orange) are separable again (dashed line) in *rsc8*-depleted strains. (D) The phenomenon can be better understood when assessing the fPCs, particularly the second. For WT small genes, the first two nucleosomes are still identifiable (+1 turquoise arrows), whereas the superposition of genes with *≤* 1000 bp make it impossible to represent the profile thereafter (+4 blue arrows, +6 orange arrows). (E) The phasing for small genes is re-established in *rsc8* conditions, as indicated by the second fPC (+4 blue arrows, +6 orange arrows). It is of note that the phasing also changes for the +1 nucleosome (+1 turquoise arrows), and the NDR can be seemingly not maintained. In Figs (D) and (E), the dashed black line as well as the solid lines in magenta and green indicate the mean, a positive contribution of the fPC, and a negative contribution, respectively.

However, it was highly surprising that the neat separation between the two clusters fully vanished for small genes (Fig 2(B), for replicate B SFig A.2 (E)). Almost all data points belong to the same group, although both are present. We want to remind that we first established the two Pearson clusters with respect to all coding regions and only *thereafter* separated small from large genes (in the following *all-gene* clusters). Moreover, the fPCs for small genes include overlapping positioning inside and outside the gene body due to their varying sizes. The fact that the clusters are not separable implies that coordinated nucleosome phasing disappears after the Transcription Termination Site (TTS), and we hypothesised that the arrangement is strictly limited to the gene body. Indeed, the second small-gene fPC indicates well-defined positioning only for up to the +2 nucleosome (*≈*300 bp), and the function loses quickly its frequent wave-like shape thereafter (Fig 2(D)). As the barrier model proposes well-defined phasing with respect to promoter-dependent properties, we analysed a possible impact of the downstream NDR by considering exclusively very large genes (*≥* 3000 bp). We did not observe any different behaviour, and the boundary was clearly visible (SFigs A.2(G, H)). We therefore surmise that the effect of the downstream NDR on phasing in the gene body might be negligible, and it is predominantly determined by the +1 position. This outcome is in line with the barrier model. To verify our hypothesis of gene-size dependent nucleosome phasing, we divided the regions into small and large genes *before* performing the Pearson clustering. When considering exclusively small genes, the two Pearson groups become linearly separable again, which is—in accordance with our hypothesis—predominantly determined by the size (SFig A.4). This shows that the nucleosome arrangement is strictly limited to the gene body.

The data produced by [12, 18] contain two replicates for *chd1*Δ, *isw1*Δ, and *isw2*Δ cells as well as *rsc8*-depleted strains, together with their combinations as double, triple, and quadruple mutants. In order to analyse how gene-size dependent nucleosome phasing alters in varying contexts, we compared the small-gene fPCs in mutant and WT conditions. Surprisingly, the separation of the two *all-gene* clusters was clearly visible for small coding regions in *rsc8*-depleted strains (Fig 2(C)). Indeed, the average MNase-seq profile exhibits phased peaks along the entire 1000 bp-window (Fig 2(E)), and nucleosome positioning continued outside the gene boundaries. The linear separability can be found in almost all mutants which are depleted of Rsc8 (SFig A.5), with the sole exception of Rsc8-depleted *chd1*Δ strains (SFig A.6(A, B)). Here, the fPCs for small genes resemble the small-gene fPCs in WT conditions, indicating that the gene-specific boundaries for nucleosome phasing can be re-established (SFig A.6(C, D)). Consequently, we hypothesise that Chd1 and Rsc8 have partially antagonistic roles for maintaining chromatin organisation that distinguishes transcribed from non-transcribed regions. Taken together, this analysis exhibits strictly constrained and Rsc8-mediated nucleosome organisation within coding regions.

### Nucleosome Phasing Changes In Remodeler Mutants

We were particularly interested in how nucleosome remodeler complexes affect coordinated phasing. To remove any gene size-dependent bias from the further downstream analysis, we applied the Pearson clustering to exclusively large genes (*>* 1000 bp) for all strains and determined their two major fPCs (SFigs A.2(C, D); A.3). We can confirm that all created groups were again significant (outside the 95% PI of a gamma distribution), with the exception of *isw1*Δ*rsc8* replicate *B* (SFig A.7). We consequently removed this value from the analysis. We determined the separating boundary for all strains by using a linear Support Vector Machine (SVM). As aforementioned, the boundary is vaguely sloped when only considering large genes in WT conditions, and the two available replicates differed slightly. The observed deviation between replicates was used as a reference for the anticipated variability in the data. Indeed, we observed notable differences between the mutants, showing that fPCA is sufficiently sensitive to capture strain-dependent consequences (SFigs A.8 and A.9 for replicate A and B, respectively). It is noteworthy, however, that the boundary between the two Pearson coefficient clusters remains largely intact, despite some of the mutants exhibiting a larger overlap between these groups. This result shows that—whilst some gene deletions caused reduced gene-wide phasing along the gene body—no mutation could completely remove the importance of coordinated positioning in an array for genes longer than 1000 bp.

We identified 5 mutants—namely *chd1*Δ, *isw2*Δ*chd1*Δ, *rsc8*-depleted *chd1*Δ, *isw1*Δ*isw2*Δ, and *rsc8*-depleted *isw2*Δ*chd1*Δ—that evoked notable changes considering the experimental variability by using Eq 6 (Fig 3). For a correct interpretation of the results, it is crucial to highlight that this does not imply that other mutants had no effect on the nucleosome profile. In fact, other gene deletions that were not marked as being significant resulted also in visibly altered MNase-seq signals. However, our measurement suggests that coordinated phasing is notably altered in these 5 mutants, and the functional deviation from a mean profile changes with respect to the WT. Other gene deletions can have other impacts that do not disrupt the collective positioning. All measurements are given in Table 1.

**Figure 3:**
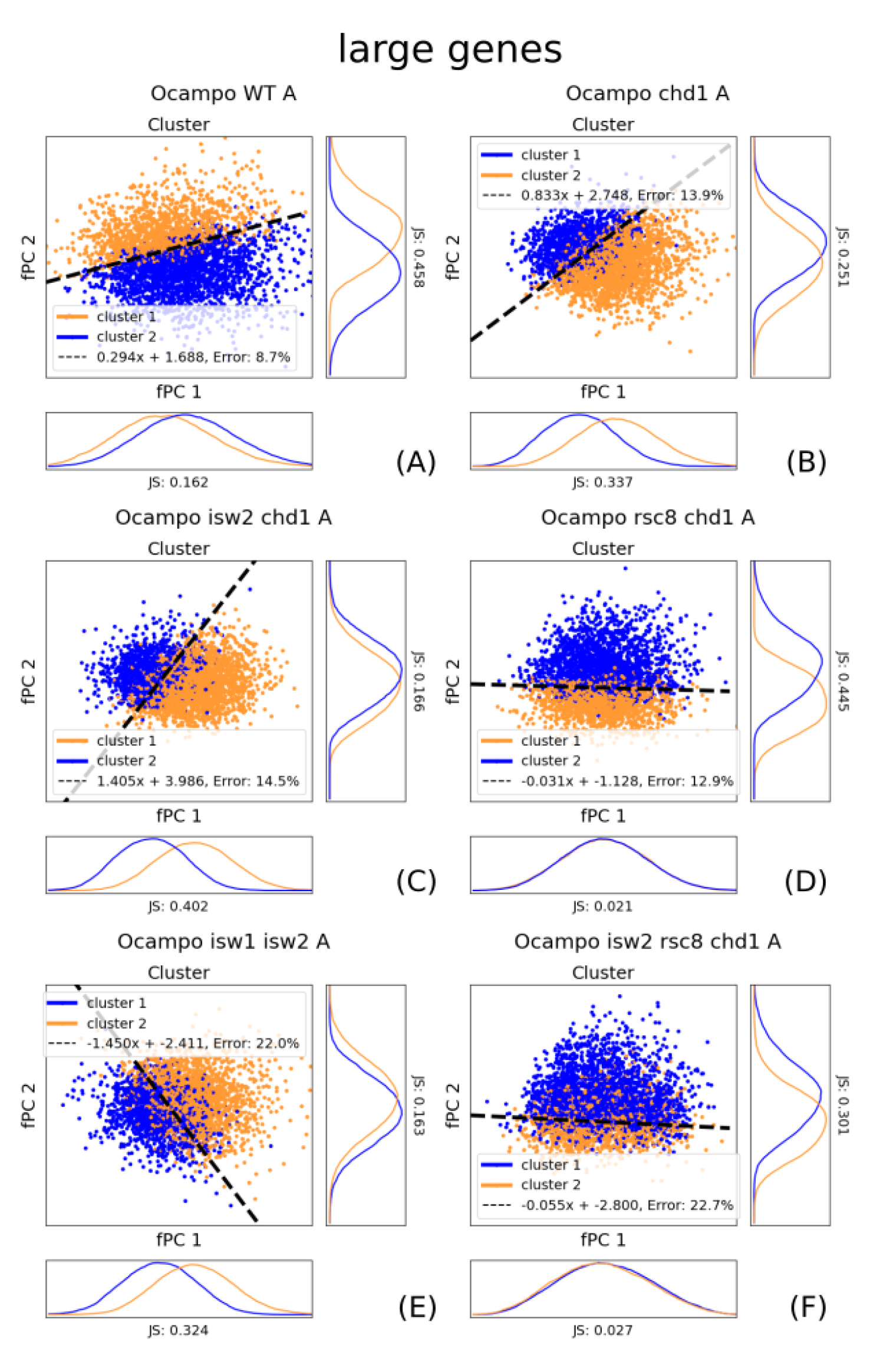
The linear separability between the Pearson clusters changes in mutants. The figure shows the fPC scores *ζ* of all conditions where the boundary slope changed notably are coloured with respect to the Pearson clustering using only large genes (*≥* 1000 bp). Blue and orange indicate each one group, the dashed line symbolises the best linear separation using a SVM. The x-axis represents the score of the first fPC *ζ*_1_, the y-axis gives the score for the second fPC *ζ*_2_. All axes are scaled to the same size; shapes are therefore comparable.

**Table 1:** SVM boundary slopes for both replicates. The first two rows give the boundary slope for replicate A and B, an *µ* is the mean slope for both. The **s** value represents our significance measurement defined in Eq 6. Noteworthy oundary slope are marked in green (bold), all others are red. The *B* replicate of *isw1*Δ*rsc8* was not significant, and moved from the analysis (crossed out). The WT is per definition equal 0.

|  | WT | chd1 | isw1 | isw2 | rsc8 | isw1/chd1 | isw2/chd1 | chd1/rsc8 |
| --- | --- | --- | --- | --- | --- | --- | --- | --- |
| **A** | 0.299 | 0.834 | 0.117 | 0.133 | 0.377 | 0.213 | 1.406 | 0.031 |
| **B** | 0.055 | 0.48 | 0.329 | 0.038 | 0.08 | 0.283 | 0.538 | 0.074 |
| **Mean $\mu$** | 0.177 | 0.657 | 0.223 | 0.0855 | 0.2285 | 0.248 | 0.972 | 0.0525 |
| **s** | 0 | **2.6674** | 0.0409 | 0.3612 | 0.0366 | 0.2951 | **2.9842** | **1.4773** |
|  | isw1/isw2 | isw1/rsc8 | isw2/rsc8 | isw1/isw2<br>chd1 | isw1/chd1<br>rsc8 | isw2/chd1<br>rsc8 | isw1/isw2<br>rsc8 | isw1/isw2<br>chd1/rsc8 |
| **A** | 1.452 | 0.347 | 0.153 | 0.112 | 0.216 | 0.057 | 0.466 | 0.066 |
| **B** | 1.074 | 0.072 | 0.567 | 0.295 | 0.207 | 0.068 | 0.245 | 0.174 |
| **Mean $\mu$** | 1.263 | 0.2095 | 0.36 | 0.2035 | 0.2115 | 0.0625 | 0.3555 | 0.12 |
| **s** | **12.7873** | 0.0157 | 0.3315 | 0.0157 | 0.5420 | **4.8846** | 0.5909 | 0.1233 |

Most single mutants had only a small or negligible effect on the global phasing along transcribed regions, with the exception of *chd1*Δ (Fig 3(B)). The gene deletion exhibited a noteworthy change in the slope. This suggests that the functional composition of the signal differs to the WT. Indeed, the signal strength decreases more quickly along the gene body in *chd1*Δ mutants (Fig 4(B)). Interestingly, the second fPC shows only a clear phasing of the +1 nucleosome, whereas it quickly abates, with the strongest effect on the +2. Consequently, the +1 position remains largely unaffected. Chd1 is responsible for nucleosome spacing along genes and is particularly involved in maintaining chromatin integrity during Pol II elongation. It is therefore intuitive that the *chd1* -deletion influences phasing within the gene body. This outcome shows the clear effect of chromatin maintenance by Chd1 after the +1 nucleosome.

**Figure 4:**
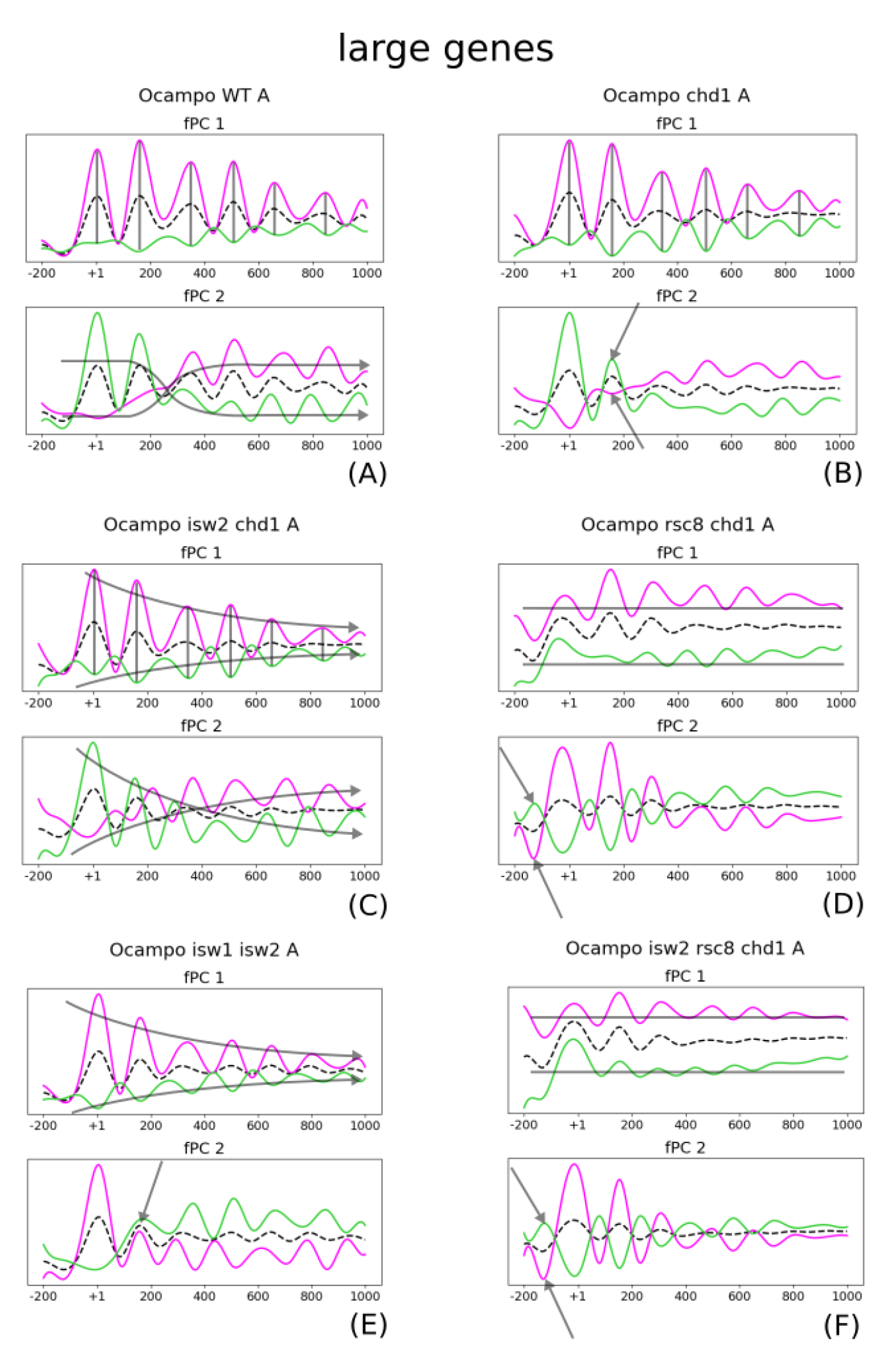
fPCA captures position specific gene-deletion effects along the entire gene on a global scale. All here presented mutants changed significantly the slope between the two Pearson correlation clusters. The lines give the mean (dashed black), a positive contribution to the mean (magenta), and a negative contribution (green), respectively. Changes that affected the amplitude are indicated with transparent black lines. Effects on phasing and single nucleosomes are emphasised with transparent black arrows. (A) The WT fPCs indicate position-dependent amplitude (fPC 1) and general changes along the nucleosome array (fPC 2). (B) *chd1*Δ changes particularly with respect to the second fPC, and the profile abates more quickly. This is particularly evident on the weak +2 positioning (arrows). (C) *isw2*Δ*chd1*Δ exhibits coupling of amplitude (transparent black lines) and gene-wide organisation (transparent black arrows). The first fPC influences particularly position-specific signal strength. Both fPCs are seemingly implicated in phasing, as the first fPC amplitude decreases as a function of distance. (D) and (F) display almost identical effects, indicating that the *isw2*Δ does not impact the observation. Most interestingly, NDR maintenance is disrupted, which is indicated by the new peak that occurs next to the +1. (E) The double mutant *isw1*Δ*isw2*Δ shows disharmonious phasing, which is particularly evident by weak positioning of the +2 nucleosome (arrow).

The double mutant *isw2*Δ*chd1*Δ exhibited also a noteworthy shift of the separating boundary (Fig 3(C)), yet with very different results to the *chd1*Δ single mutant. *isw2*Δ*chd1*Δ preserves largely spacing (Fig 4(C)), however strongly entangles the first and second fPC for separating the Pearson clusters. Positions remain organised along the entire transcribed region, but the signal amplitude per peak and over the entire profile are important in both fPCs. This suggests that nucleosome presence is affected rather than their positioning in an array along the DNA. Surprisingly, the *rsc8*-depleted *chd1*Δ decreases the slope tilting (instead of accentuating it), therefore making coordinated phasing almost exclusively represented by the second fPC (Fig 3(D)). This can be better understood when analysing their respective effects (Fig 4(D)). The first fPC solely explains average signal amplitude and contains almost no information about coordinated positioning. Interestingly, the second fPC indicates that the NDR before the +1 cannot be maintained, which is in line with other studies [12, 39]. It should be noted that not all double mutants that include *chd1*Δ show a similarly notable impact as the single mutant. This could possibly mean that these double mutants have opposing effects, although it is difficult to give a clear indication with the variation between only two replicates. We found a mixed picture for *isw1*Δ*isw2*Δ. The strongly sloped boundary suggests that linear correlation is functionally coupled with another behaviour (Fig 4(E)). In fact, the effect of the second fPC hints that the positioning of the +2 is strongly impacted, and following positioning becomes inharmonious (Fig 4(E)). The +1 is kept well positioned. Taken together, these results show that double mutants can have varying and non-linear effects.

Among the triple and quadruple mutants, the only one that changed notably the clustering boundary is *isw2*Δ*chd1*Δ*rsc8* (Fig 3(F)). Once again, tilting is decreased. The effect is almost identical to the *chd1*Δ*rsc8* mutant, suggesting that *isw2*Δ does not have a strong effect on the phenomenon (Fig 4(F)). However, it should be mentioned that the variability between the two replicates is considerably large, as the two clusters can be only neatly separated in replicate *B*, whereas replicate *A* exhibits a great overlap. Moreover, mutants with more than two gene deletions were largely deprived from clear nucleosome peaks, and a straightforward interpretation of the Pearson correlation could be difficult. The results for these strains should be taken with a pinch of salt.

Taken together, these outcomes show that remodeler mutants have varying effects on phased nucleosome localisation. Whilst most mutations do not notably change the separation of the two Pearson coefficient clusters with respect to the WT, we identified 5 mutants that exhibited a strong effect on phasing. Interestingly, most of them include *chd1*Δ, which indicates an important role of Chd1 for local nucleosome arrangement within the gene body. FPCA permits the clear and position-specific quantification of the induced impact among varying strains.

### Pol II Presence Correlates With Nucleosome Organisation in *chd1***Δ Mutants**

In order to assess an interdependence of nucleosome organisation with other genomic properties, we compared the two Pearson clusters to Pol II levels, Sth1 occupation, AT ratio over the entire gene, as well as upstream NDR length and orientation of the upstream NDR (i.e. tandem or divergent). We also included Mediator presence as a large protein complex with transient interactions predominantly at the NDR. All of these factors were clustered into two equally-sized groups where possible. For example, Pol II presence along the gene was evenly separated into transcribed regions with high and low Pol II occupancy. Interdependence was measured by training a simple neural network with no hidden neurons using Hebbian learning [40]. Consequently, we assessed which nuclear groups (e.g. high or low Pol II presence) corresponded to which Pearson clusters. The initial *k*-mean clustering did not impose a constraint on the group size, and they could therefore differ in the number of genes they contained. To remove any prediction bias, we forced the clusters to be of the same size. Genes in the larger group with closest Pearson coefficient to all distributions in the smaller group changed the cluster. We found the analysis for WT conditions non-conclusive, and correlations varied between *A* and *B* replicate (SFig A.10(top)). Whilst *A* was slightly correlated with the AT sequence content (Figs 5(A) and (B)), this trend disappeared for *B*, and it might in both replicates rather correspond to the fPC orthogonal to the cluster boundary (SFig A.11). Overall, we were surprised that none of the investigated properties could indicate a clear interdependence with nucleosome phasing in WT (Fig 5(A)).

**Figure 5:**
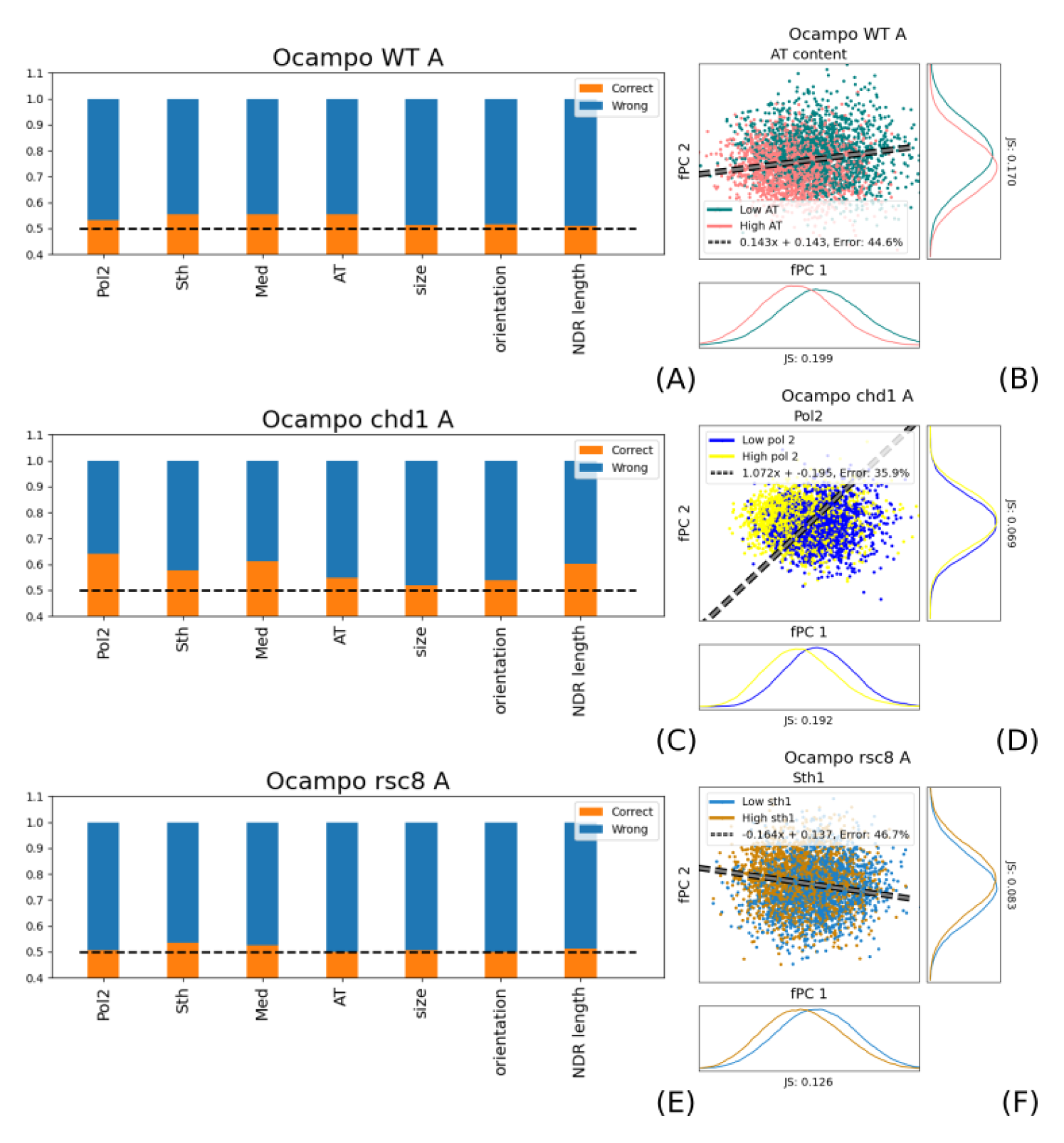
Remodeler deletions have varying effects on the interdependence with other genomic properties. The orange bars in Figs (A), (C), and (D) show the ratio of correct predictions, and blue bars are wrong guesses. As we distinguish between two clusters, the dashed black line at 0.5 indicates random guessing. The dashed grey line with black edging in Figs (B), (D), and (F) display the linear boundary for the Pearson clusters. (A) and (B): WT conditions are seemingly correlated with the sequence composition. However, the results are different for the *B* replicate, and therefore non-conclusive. All possible correlations are surprisingly low. (C) and (D): *chd1*Δ mutants increase particularly their dependence on Pol II and other transcription-coupled properties, such as Mediator presence. Surprisingly, the mutant showed also an increased interdependence on NDR length. (E) and (F): despite the Rsc8-mediated gene limits, there is no correlation with coordinated nucleosome phasing and the size of transcribed regions or NDR length. Although Sth1 indicates a slightly increased interdependence, this cannot be confirmed when plotting the groups with respect to the Pearson clusters. This is in line with our hypothesis that positioning in different regions interfere, and therefore, nucleosome localisation become increasingly independent from region-specific factors.

The correlation between positioning and other nuclear properties changed among the mutants (SFig A.10). The effect is particularly clear for *chd1*Δ (Fig 5(C)), as there is a strong interdependence between phasing and Pol II (Fig 5(D)), Mediator presence, and NDR size (SFig A.10). As aforementioned, Chd1 maintains, among others, chromatin integrity during Pol II elongation. The correlation is therefore in line with our previous conclusions and the function of Chd1. The established link between Pol II presence and nucleosome organisation remains conserved in all strains with a Chd1 gene deletion, except *isw1*Δ*chd1*Δ*rsc8*. This is similarly true for the correlation with Mediator occupancy and NDR length. There was also a slight correlation to Sth1 and AT ratio in cells containing *chd1*Δ, which were, despite being weak, still notably stronger than in WT. The results are in agreement with the effects of Chd1 on chromatin maintenance during gene expression.

Due to the Rsc8-mediated nucleosome organisation within transcribed regions, we were wondering whether there is an increased interdependence to NDR length or gene size. We can report that there is no correlation with NDR size in any *rsc8*-depleted strain (Fig 5(E)). This is in line with our hypothesis that Rsc8 decouples processes at different genes. However, *rsc8* mutant cells exhibited a slightly increased correlation with Sth1. By looking at the separation with respect to the Pearson cluster boundary, we find that there is no noticeable impact (Fig 5(F)). The results indicate that any correlation with region-specific properties is lost, which is likely due to the interference of nucleosome positioning of various regions.

*isw1*Δ single mutant did exhibit only a slightly increased correlation with Pol II, Sth1 and Mediator presence as well as AT ratio. *isw2*Δ might as well show a weak correlation with Pol II occupation. Their double mutant differ strongly between replicate A and B, and it is therefore difficult to tell whether transcription-related factors influence nucleosome phasing in the strain. However, none of them indicate any strong interaction, suggesting that—on a global scale—these effects might be negligible in comparison to the WT (SFig A.10).

Interestingly, the *rsc8*-depleted *isw2*Δ indicated a strong correlation with Sth1 and Mediator presence as well as NDR length. The effect was observable in almost all strains that contained the double mutation with the exception of the quadruple mutant (SFig A.10). Taken together, this could indicate an impact along the gene body and the promoter region in strains that contain *rsc8*-depleted *isw2*Δ.

Surprisingly, combining two factors together (e.g. Pol II presence and AT ratio) to predict Pearson clustering did not increase accuracy. Instead, one factor dominated the correlation measurement, e.g. Pol II presence for *chd1*Δ strains. This could possibly suggest that—despite several factors showing increased interdependence—they can be reduced to a main influencing factor (which is not necessarily one of the tested properties).

Taken together, the results indicate a strong interdependence between local genomic properties—such as presence of large protein complexes or NDR length—and cell strains containing *chd1*Δ. This supports our hypothesis of Chd1 being responsible for local nucleosome coordination.

## Discussion

In this work, we analyse far-reaching nucleosomal interdependence within the gene body in WT and chromatin remodeler-deficient strains using fPCA, an analysis framework for functional data. Although fPCA is well established in the assessment of time series, it has not been previously used to understand location-specific nucleosome profiles on a global scale. By determining the two major fPCs for all transcribed regions, we quantified the effect of coordinated phasing with respect to the +1 position. This reveals the impact of different gene deletions of chromatin remodelers on nucleosome arrangement within the gene body. Our analysis reveals strict Rsc8-defined boundaries for nucleosome positioning along the gene. We determined two significant Pearson clusters with distinct phasing, which we compared with other nuclear properties—such as Pol II presence and NDR maintenance—and sequence-dependent characteristics. None of the commonly supposed influencing factors can easily explain coordinated nucleosome positioning in WT conditions. However, correlation between tested properties and phasing increases with some gene mutations. In the following, we critically discuss the results and their significance.

We classified genes according to their Pearson coefficients by applying a *k*-mean clustering approach. *k*-mean was repeated over several random initialisations, therefore removing any prior bias. We used a silhouette criterion value to determine the best number of clusters, which was shown to be 2. The validation using the silhouette criterion together with the shape-independent JS distance over 500 random clusters proved their significance. We acknowledge the fact that 500 random partitions for over *≈*5000 transcribed regions is still comparatively low. However, as we approximate the distribution of JS distances over random clustering with a gamma distribution, we made our estimates independent of the actual number of samples. Gamma distributions are commonly used to represent unimodal strictly positive random variables, and it is therefore a sensible choice for JS distances over random partitions of the same data set.

FPCA assumes a mean behaviour over the entire data set and characterises each data point with respect to their deviance from that mean (see Methods). The results therefore depend on the entire considered data set. Indeed, we find different results when including all genes or exclusively transcribed regions *>*1000 bp. However, these differences are not strong. Moreover, any possible bias was excluded by removing genes smaller than 1000 bp from a subsequent analysis. Due to the abundant and well-positioned nature of nucleosomes within the gene body in *Saccharomyces cerevisiae*, we find it justified to presume an average nucleosome distribution describing their wave-like profile. We found that the two Pearson correlation clusters could be neatly separated by the fPC scores *ζ_i_*, *i ∈ {*1, 2*}*. Consequently, the groups remain preserved in an independent evaluation of the profile. This further supports our claim that deviation from an average profile is a reasonable representation of nucleosome positioning. However, whilst linear-correlation measurements are limited to quantifying the average similarity, fPCA allows characterising location-specific differences and in which way gene deletions affect phasing from an average. Therefore, fPCA is in line with previous ways of analysing nucleosome distributions using Pearson [34] or autocorrelation [33], but significantly widens the assessment scope.

The analysis can clearly distinguish between mutant-specific dynamics on phasing. All mutants preserved the information of coordinated nucleosome arrangement in their first two fPCs, and the Pearson clusters could be separated by a neat line. Consequently, none of the chromatin remodeler gene deletions caused random positioning. Some mutants, however, showed an increased overlap between the two groups, which indicates increased independence between individual nucleosome locations, and positioning might be more random. Nonetheless, most strains did not alter notably their functional composition, and a linear separation of the Pearson clusters using the deviance from the mean did not change with respect to WT strains. Although they can nevertheless have a drastic impact on the mean itself, coordination along the genes remains preserved in a similar way. Due to the focus on the study on coordinated nucleosome arrangement along transcribed regions, we did not consider them as having notably changed their coordinated phasing.

Gene mutations of chromatin remodelers have been analysed previously in detail, including their influence on phasing [12–14, 18], NDR maintenance [41], and gene transcription [13]. RSC is the only essential chromatin remodeler complex in *Saccharomyces cerevisiae* [42], and it has been particularly associated with positioning of the +1 and −1 nucleosomes [12, 41, 43]. This mechanism has been proposed to be conserved among various yeast species [11]. It has also been reported that RSC regulates expression of Pol II and Pol III-transcribed genes [13, 44, 45]. Moreover, it has been found to impact Pol II elongation and transcription termination [12]. All of these results imply that RSC is to some extent involved in limiting the transcribed region. However, this has been predominantly quantified with respect to changes at the core promoter. To our knowledge, a potential role for Rsc8 to decouple nucleosome phasing in independent genes has not been suggested. The presented functional analysis of MNase-seq profiles in *rsc8*-depleted strains clearly indicates a coordinated nucleosome arrangement that exceeds the limits of transcribed areas. This is further supported by our finding that correlation with other nuclear and sequence-dependent factors decreases. Furthermore, mutants that were *rsc8* depleted decreased notably the boundary slope between the two clusters, indicating that coordinated positioning becomes increasingly independent of other functional components. The strictly limited and Rsc8-mediated phasing barrier could have further implications for other processes—such as transcription—as nucleosome placing in one gene influences its neighbouring regions. The notion of gene-interfering positioning has been also proposed by [14]. The study shows that RSC could act as a bidirectional barrier, influencing upstream and downstream regions. Interestingly, they found that interference also plays a crucial role in WT strains, and that the same phenomenon remains preserved in *rsc8*-depleted cells. However, our fPCA reveals that the limiting role of the RSC remodeler complex is crucial in WT conditions, and that this behaviour is significantly altered when Rsc8 is depleted. Taking this into account, Rsc8 should fulfill the role of disentangling gene-related processes in WT strains, and it therefore allows for a flexible and uncorrelated transcriptional program. Indeed, *rsc8*-depleted cells exhibit significantly altered Pol II profiles [10, 12], which is in accordance with our hypothesis. We propose that the RSC chromatin remodeler globally disentangles nucleosome phasing, and it therefore plays a substantial role in long-range positioning.

Interestingly, our results indicate that positioning limited to the gene body can be re-established in *rsc8*-depleted *chd1*Δ mutants. We hypothesise that they have antagonistic effects in establishing gene size-dependent barriers for nucleosome arrangement. Indeed, it was reported that that Rsc8 and Chd1 have opposing effects for Pol II termination. *rsc8*-depleted cells exhibit inhibition of Pol II dissociation at the TTS, whereas the double mutant *isw1*Δ*chd1*Δ increases release frequency, with seemingly *chd1*Δ dominating this effect [12]. The authors propose that this is related to the close packaging of nucleosomes at the TTS. Our outcomes suggest that they might have antagonistic effects in chromatin organisation that differs between transcribed and non-transcribed regions.

We found that *chd1*Δ mutants had a strong impact on coordinated positioning within the gene body. Indeed, Chd1 has been, among others, characterised with respect to its role in maintaining chromatin integrity during Pol II transcription [16, 46, 47], and it associates to both promoters and transcribed regions [48]. This is in line with our finding that correlation with Pol II presence and occupancy of Mediator increases in Chd1-deficient strains. With the exception of *isw1*Δ*isw2*Δ, all other noteworthy changes included deletion of *chd*, further emphasising its role for chromatin organisation within the gene. However, not all *chd1*Δ-containing mutants exhibit a notable effect. This can have various reasons, including experimental variability. However, particularly the mutant *chd1*Δ*isw1*Δ*isw2*Δ could indicate an interacting behaviour of the remodelers. Indeed, Chd1 has been reported to cooperate [16] as well as antagonise Isw1 [18], and therefore could have different effects depending on the context. With this being said, the behaviour of the triple mutant *isw1*Δ*isw2*Δ*chd1*Δ is particularly interesting, as *chd1*Δ and *isw1*Δ*isw2*Δ each individually affect coordinated phasing, but not their triple mutant. This could suggest an antagonistic behaviour on nucleosome coordination. As Chd1 is highly conserved in all eukaryotes [49], this result could have consequences beyond *Saccharomyces cerevisiae*.

Analysing the MNase-seq data using fPCA allowed us to obtain a different view on the functionality of various remodelers to maintain chromatin organisation. We propose the following mechanism (Fig 6). The RSC remodeler complex is essential for allowing independent phasing in each single gene. It plays therefore a pivotal role in maintaining the barrier with respect to which nucleosome positioning is coordinated. This permits the decoupling of gene-specific processes such as transcription. Depletion of Rsc8 leads to the interference of different genomic regions, which therefore alters sequence accessibility on a global scale. Indeed, it has been reported that gene expression is dramatically changed in *rsc8* mutants [10, 45]. Chd1, on the other hand, maintains chromatin integrity during transcription [16, 46, 47], and it influences nucleosome phasing locally to permit Pol II-mediated expression. *chd1*Δ strains make positioning dependent on Pol II presence. Consequently, whilst RSC plays a global role, Chd1 is important for local nucleosome organisation.

**Figure 6:**
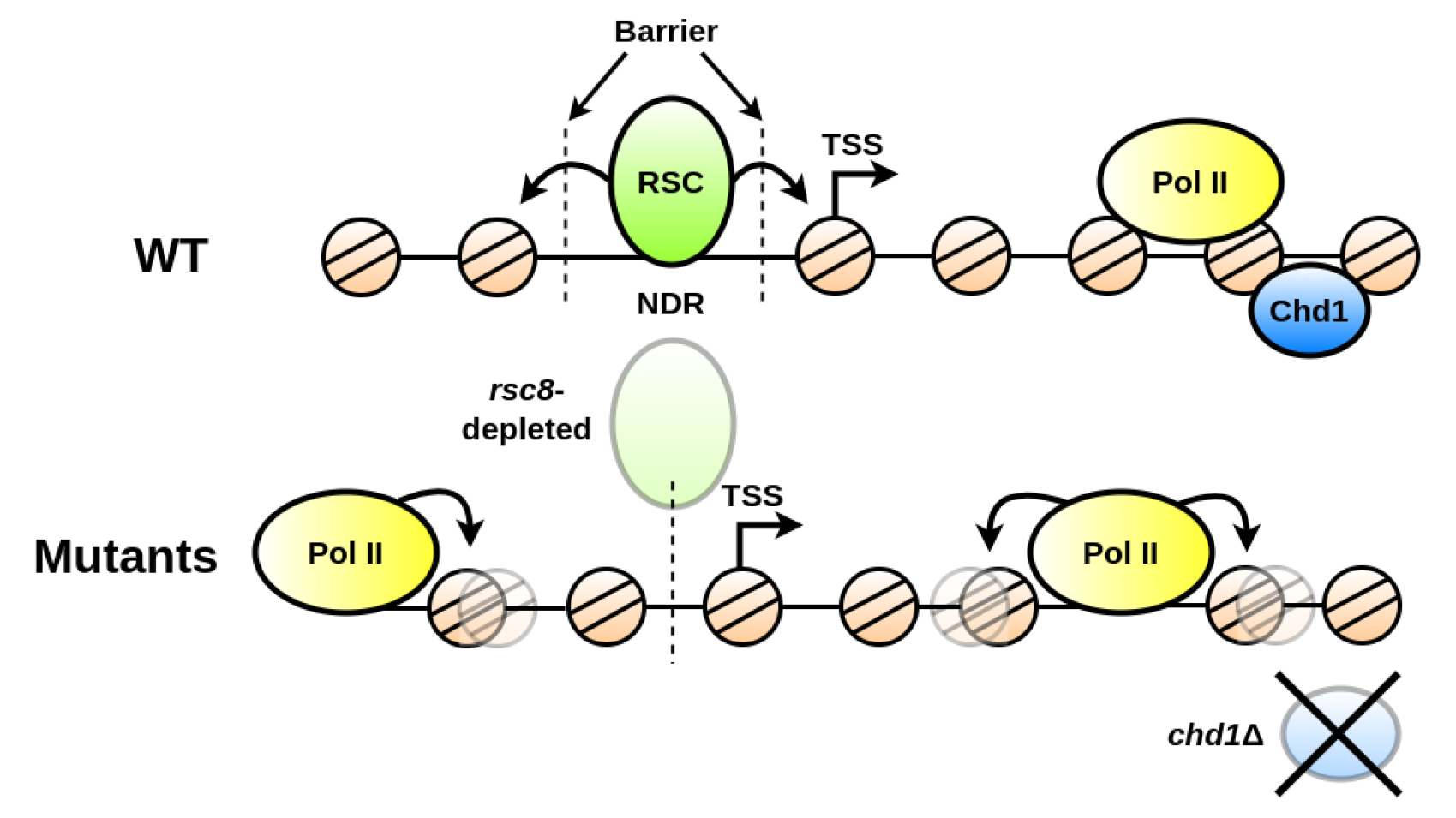
Chromatin remodelers maintain nucleosome organisation on a local and far-reaching scale. Top: RSC (green ellipse) establishes independent nucleosome phasing on each gene (two vertical dashed lines) by maintaining the NDR through positioning the +1 (cornered arrow) and −1 nucleosome. The ATP-dependent positioning is symbolised by black arrows pointing away from RSC. The local remodeling effect of Chd1 (blue ellipse) allows chromatin arrangement independent of Pol II transcription (yellow ellipse). Bottom: in *rsc8* strains, the NDR cannot be maintained anymore, and phasing in and outside a gene interfere with each other (single dashed line). We propose that this should equally lead to an increased interdependence of other nuclear processes such as transcription. If *chd1* is deleted, nucleosome arrangement is more sensitive to the presence of other large complexes, such as Pol II. During transcription, Pol II is affecting the local positioning (black arrows from Pol II).

## Methods

### Data Treatment

MNase sequencing reads were taken from [18] and [12] (GEO accession numbers GSE69400 and GSE73428, respectively) and treated as in our previous study [50]. To be precise, reads from Fastq files were trimmed with trim galore (v0.6.5) [51] and cutadapt (v3.1) [52]. Subsequently, they were mapped on the *Saccharomyces cerevisiae* genome (University of California at Santa Cruz [UCSC] version sacCer3) using bowtie2 (v2.3.4.3) [53]. Files were converted with samtools (v1.9) [54] and deeptools (v3.5.0) [55]. Read counts were normalised in Reads Per Million (RPM) of mapped reads. We used the option --MNase of bamCoverage so that only the mononucleosome fragments were kept. This means that fragments shorter than 130 bp and longer than 200 bp were removed from analysis. Mediator and Sth1 ChIP-seq were taken from [50] (ArrayExpress accession number E-MTAB-12198). We used Pol II ChIP-seq from our previous study [56].

Following [12, 18], we retrieved positioning profiles along the coding regions 200 bp before and 1000 bp after the +1 nucleosome. Genes on the Crick strand were inverted. Consequently, all data is calibrated such that the +1 position is at 200 bp. The profile of genes for which the +1 position is known were considered as in [18].

Clustering was performed by determining the inter and intra Pearson correlation of MNase-seq distributions per gene (Fig 7(A)). To define the optimal number of *k*-mean clusters, we used the silhouette criterion measurement [57, 58]. For all analysed strains, the highest silhouette value occurs at 2 groups, suggesting that the optimal number of clusters is 2 (Fig 7(B)). Therefore data were grouped in two *k*-mean clusters (Figs 7(C, D)).

**Figure 7:**
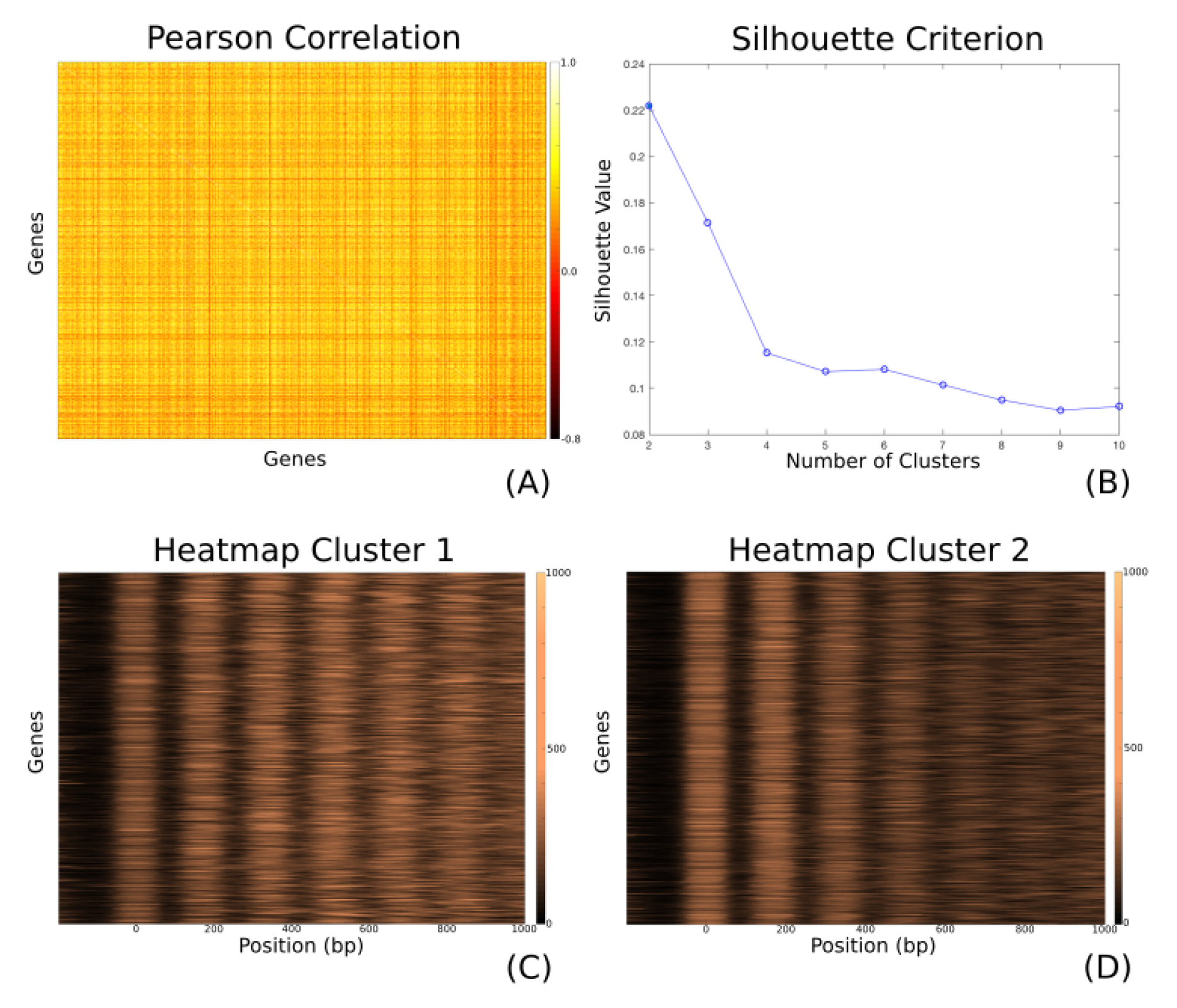
Example of grouping data to two clusters for the WT strain. (A) The heatmap shows the inter and intra Pearson correlation matrix for 5479 considered genes. Each row and column represents a gene. The color map represents the Pearson correlation coefficient. (B) Silhouette plot clearly indicates that the data can be best divided into two clusters. Although, the values are comparatively low (*≈* 0.22 for two clusters), we can find a neat separation between the groups when applying fPCA. (C) and (D) A decrease of nucleosome as a function of distance from the +1 becomes clearly visible in cluster 2 When aligning the profiles of nucleosome dyads. Each row represents a gene, and the x-axis shows the position along the coding region, with the +1 nucleosome defined to be at position 0 bp. The color code represents the population-averaged presence of a dyad to the range [0, 1000].

Cluster significance was validated by comparing the JS distance of the two determined groups with 500 random groups. However, standard significance tests are performed by comparing two distributions, instead of comparing a single value (JS distance of the Pearson clusters) to a distribution (500 random JS distances). We therefore approximated the underlying function with a gamma distribution and determined its 95% PI. If the value was outside the PI, we deemed it to be significant. Unfortunately, fits to the gamma distribution were sensitive to outliers, such that a single value far from the mode had a stronger weight in comparison to each value around it. We presumed that the distribution is best represented with the values close to the mode and removed therefore the 99% percentile of the random JS distances before fitting.

### Functional Principal Component Analysis (fPCA)

Functional clustering in a Hilbert space *H* can be achieved by fPCA. It applies—similar to PCA in Euclidean space—a functional dimensionality reduction in *H* to investigate the dominant mode in functional data. Instead of relying on values in discrete dimensions, fPCA uses a given number of basis functions (e.g. B-splines or Wavelet) to create the eigenfunction basis that accounts for most functional variation. Despite the fact that MNase-seq data is stored in a discrete array (i.e. one value per bp), we can nevertheless find a functional approximation over a range using a given choice of basis functions. It should be noted that this implicitly smooths out high frequencies in the signal. We presume that nearby values in MNase-seq data possess a strong interdependence, therefore justifying a smoothed and continuous functional representation of the high dimensional data. In this study, we apply B-splines as a basis to represent the nucleosome array (Fig 8). We use the Python library scikit-fda to determine the fPCs and the corresponding weights explaining the distribution [59]. Here, we describe briefly the underlying principles of the method.

**Figure 8:**
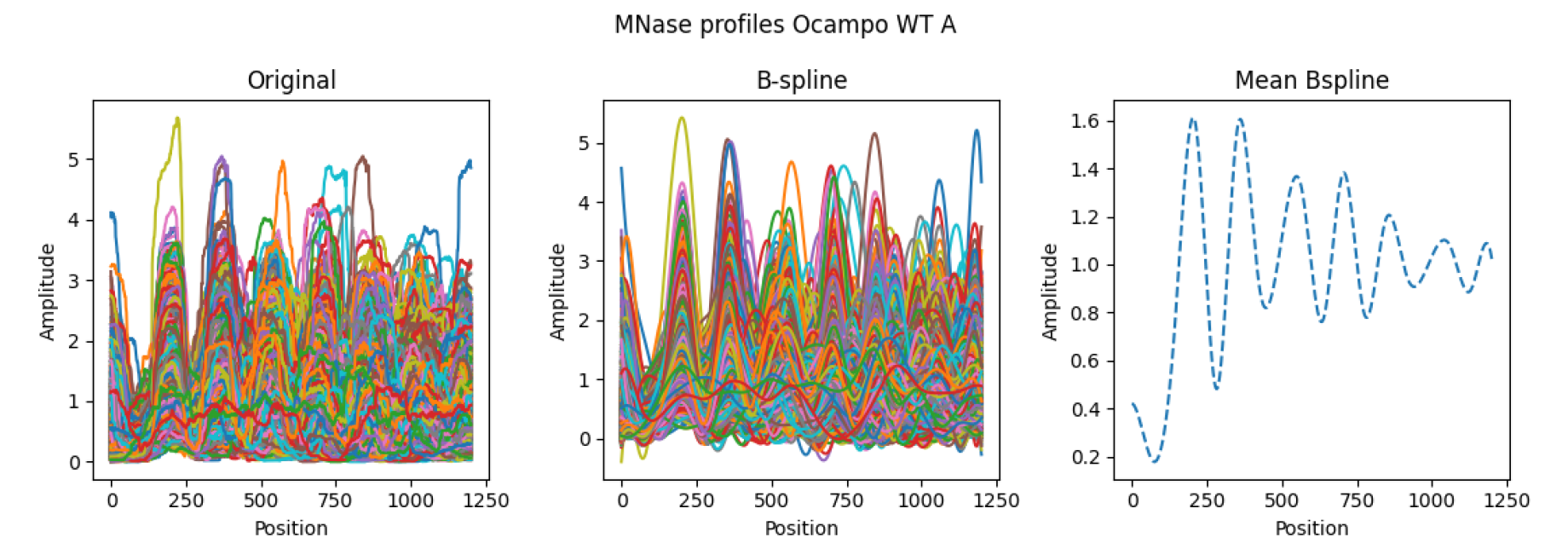
Representing the MNase data array as a composition of B-spline base functions in WT conditions. Left shows the raw data, each colour depicting one profile over a gene. Center gives the smoothed profiles after representing the data as B-splines. Right displays the average profile using the functional composition.

FPCA presumes that the functional data represents a stochastic process *X*(*t*) with expected value *µ*(*t*) = E [*X*(*t*)] and orthonormal eigenfunctions *ϕ_i_*(*t*), *i* = 1, 2…. Intuitively, *ϕ_i_*(*t*) describes the most variation in *X* orthogonal to all *ϕ_j_*, *j < i*. This allows the iterative determination of the eigenfunctions in the functional data. It should be emphasised that in this study the process is defined in space rather than describing temporal data. We follow nevertheless the convention by denoting the independent variable as *t*. By using the Kosambi–Karhunen–Loève theorem, any stochastic process can be represented as an infinite linear combination *ϕ_i_*(*t*). Consequently, we can describe the stochasticity in *X*(*t*) via

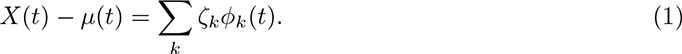

*ζ_k_* is the autocovariance operator

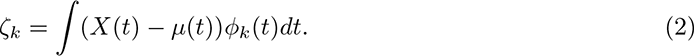

To provide some intuition, it is presumed that the entire data set can be explained via an average behaviour *µ*(*t*). Variability to *µ*(*t*) for each gene is expressed by *ϕ_k_*(*t*) together with a factor *ζ_k_*. *ζ_k_* can be loosely compared to a normal correlation measurement, i.e. *ζ_k_* increases when *ϕ_k_*(*t*) and *∫*(*X*(*t*) *− µ*(*t*)) follow the same trend. If they describe opposing behaviours—for example decreases when *∫*(*X*(*t*) *− µ*(*t*)) increases—*ζ_k_* becomes negative.

It is commonly justified to approximate Eq 1 as a finite sum

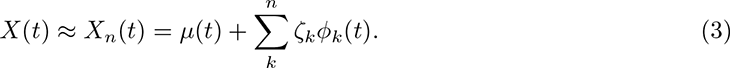

It should be noted that *ϕ_i_*(*t*), *i* = 1, 2, … is a basis of the functional space in *H*.

This understanding of the underlying process permits the application of fPCA. A smoothed representation with the basis functions (e.g. B-splines) fulfilling Eq 3 can be obtained using *L*^2^ regularisation. To reduce the dimensionality to *K*, we keep only the first *K* components (i.e.*ϕ_i_*(*t*)) that represent the dominant mode of variation in *X* by setting the first component to

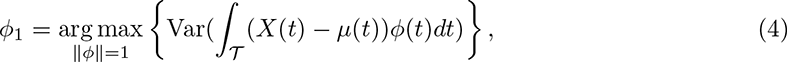

and the following *K −* 1 components to

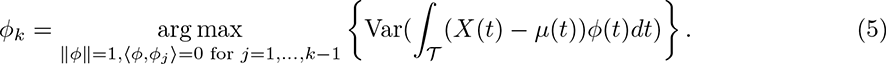

∥*ϕ*∥ is the square norm, i.e. 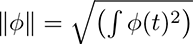. It should be emphasised that *ϕ_k_* can differ by a factor of −1 due to the square norm, and consequently, the operator *ζ_k_* (Eq. 2) can be either positive or negative depending on *ϕ_k_*. This means that the slope of the cluster-dividing boundary can bee pointing upwards or downwards and still describe the same functional composition.

We exemplified the impact of the first two fPCs to analyse the consequences on nucleosome phasing in chromatin remodeler-deficient cells (see for example Figs 1, 2, and 4). It should be noted that the fPCs were amplified to highlight their functional contribution. We set *ζ*_1_ = *ζ*_2_ = 20 in all figures that demonstrate their effects. The determined *ζ*_1,2_ *∈* [*−*20, 20] for all strains and replicates, and most of them were in fact much lower.

### Quantifying the Cluster Boundary

Long genes were linearly separable with respect to the Pearson coefficient clusters in all WT and mutant conditions. The boundary was determined using a linear SVM. We ignored the prediction error and the intercept of the linear boundary, and instead considered only the slope differences between the two replicates. As aforementioned, the sign of the slope *m* does not matter, and we consider therefore only *|m|*. To quantify the variability in the two replicates, we introduce the following measurement

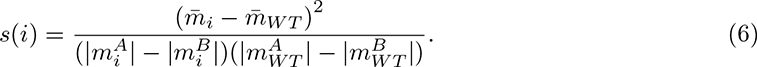

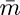 denotes the average over the absolute slopes of both replicates. We defined a change as notable when *s*(*i*) *>* 1, which implies that the mean variability between WT and mutant is larger than the variability within the replicates, i.e.

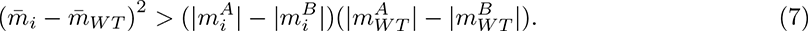

As we consider only two replicates, we restrain from using the word *significant* as much as possible and use *noteworthy* or *notable* instead.

### Measuring Interdependence Between Nucleosome Phasing and Other Nuclear Properties

In order to analyse interdependence of nucleosome positioning with other nuclear properties, we divided all factors into two equally sized cluster using the median wherever possible. For example, the half with the smaller NDRs was assigned to group −1, whereas the larger half was group 1. This split was performed after filtering for the size (i.e. large or small genes). The analysis aimed to find a correlation between nuclear factor group and Pearson cluster. To remove any bias with respect to the group size, we forced both Pearson clusters to contain the same number of genes.

We used a simple feedforward network with no hidden neurons and a single output neuron whose activation indicated the predicted Pearson cluster. The number of input neurons varied between 1 and 2, depending on whether we considered a multivariate interdependence. The group of the nuclear factor (i.e. −1 or 1) was set as input neuron activation. This was weighted and summed together with all other input values. The activation function of the output was a modified sign function, which returned 0 when negative and 1 when positive. Therefore, if the weighted sum over the input was lower than or equal to 0, the output would be 0, and 1 otherwise.

Weights were trained using a Hebbian-like learning method [40]. In order to avoid any confusion, we name Pearson cluster 0 and nuclear factor group −1 *low* cluster, whereas we define group 1 in both cases to be the *high* cluster. The weight was defined to be the average number of genes where the nuclear factor group and Pearson coefficient cluster where both *low* or both *high*; minus the average number where one of the was *low* and the other *high*. The implementation as a neural network allowed the straightforward extension to compare interdependence with several factors at the same time using the same method.

## Acknowledgments

This work was supported by Fondation ARC [PGA1 RF20170205342]; Comité Ile-de-France - La Ligue Nationale Contre le Cancer. K.A. was supported by a PhD training contact from the French Ministry of Higher Education and Research. L.Z. was supported by a PhD training contract from the CEA NUMERICS program, which has received funding from European Union’s Horizon 2020 research and innovation program under the Marie Sklodowska-Curie grant agreement No 800945.

## A Supplementary Figures

**Figure A.1:**
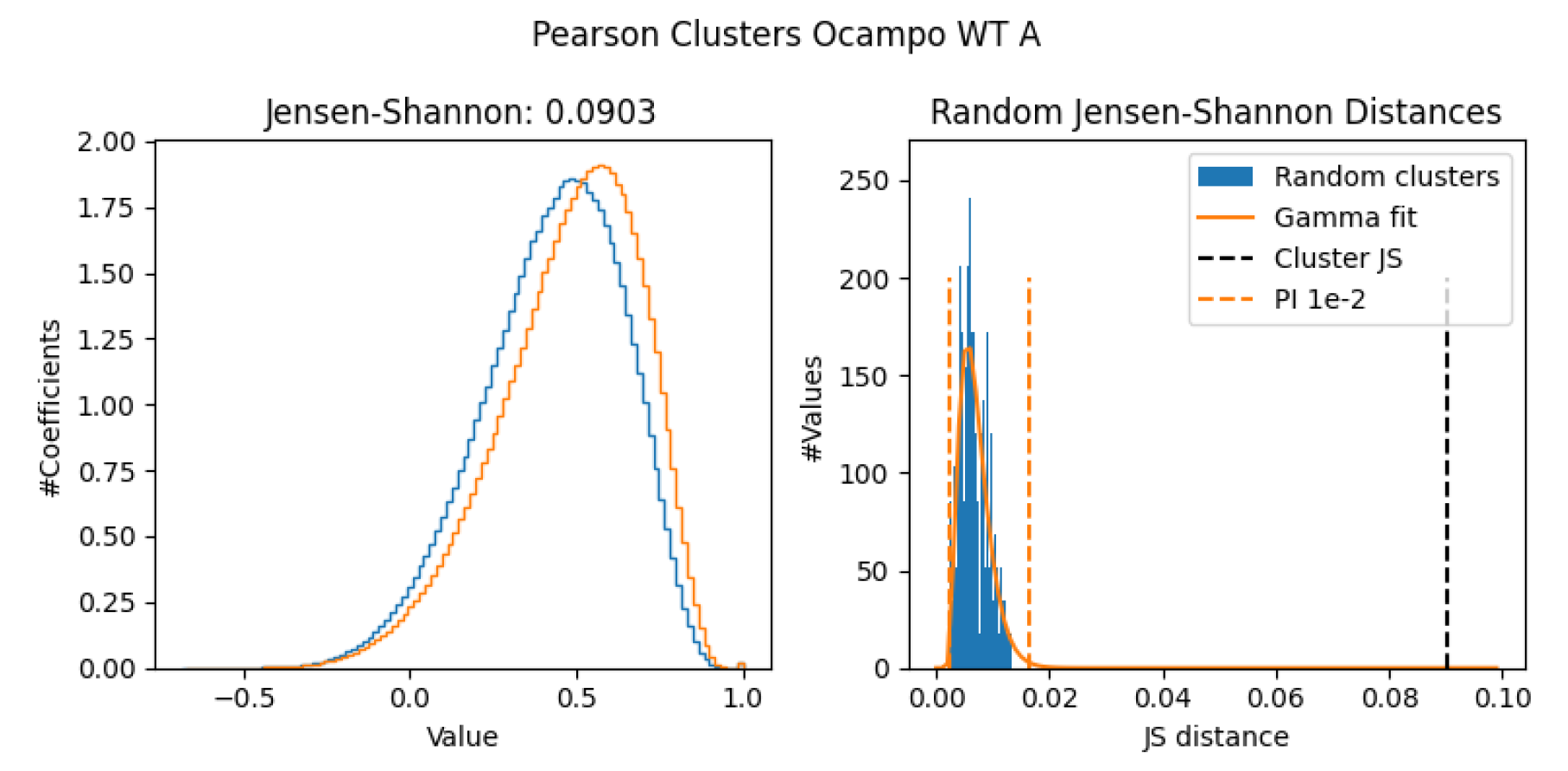
Cluster significance test for all genes. Left: The Pearson correlation coefficients for each profile (blue and orange) in a cluster to all other distributions (independent of the cluster) is seemingly very similar for both groups, as indicated by shape-independent the JS distance. Right: By measuring the JS for 500 random and mutually distinct clusters (blue bars), we can approximate the expected distance over two random groupings of the Pearson coefficients using a gamma distribution (orange solid line). Indeed, the JS between our initially determined clusters (dashed black line) is outside the 99%-PI (dashed orange lines), proving that the separation is significant.

**Figure A.2:**
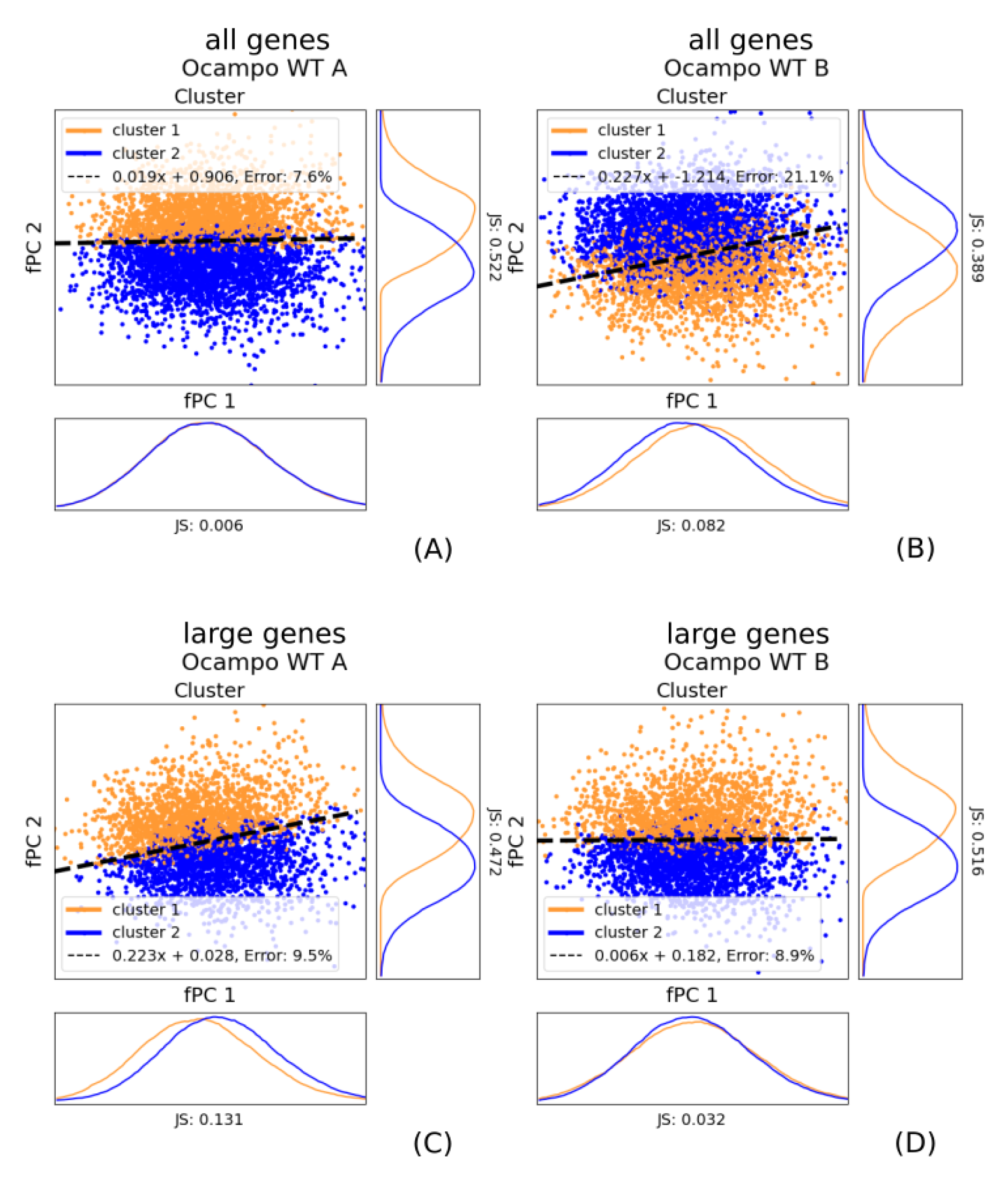
The WT fPC scores *ζ* coloured with respect to the Pearson clustering using all genes (part 1). Blue and orange indicate each one group, the dashed line symbolises the best linear separation using a SVM. The x-axis represents the score of the first fPC *ζ*1, the y-axis gives the score for the second fPC *ζ*2. All axes are scaled to the same size; shapes are therefore comparable. (A) and (B) show all genes for replicate A and B. (C) and (D) display the fPC scores after filtering for large genes (*>* 1000 bp) for replicates A and B.

**Figure A.2:**
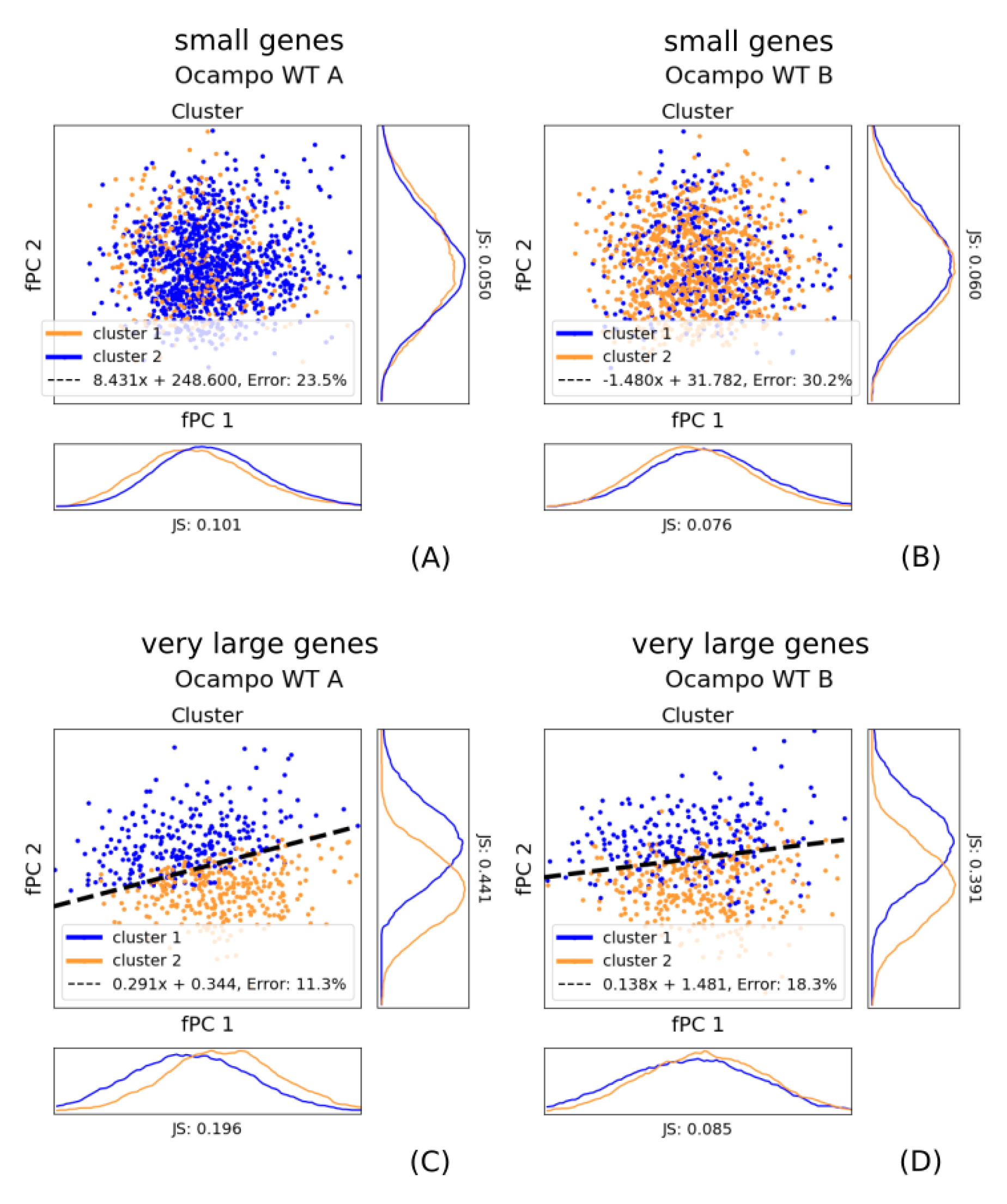
The WT fPC scores *ζ* coloured with respect to the Pearson clustering using all genes (part 2). Blue and orange indicate each one group, the dashed line symbolises the best linear separation using a SVM. The x-axis represents the score of the first fPC *ζ*1, the y-axis gives the score for the second fPC *ζ*2. All axes are scaled to the same size; shapes are therefore comparable. (A) and (B) show small genes ((*≤* 1000 bp) for replicate A and B. We removed the separating boundary because it did not reasonably divide the clusters. Nevertheless, we kept the estimated linear function in the legend to allow a comparison with other boundaries. Of particular note is the bias, which can be even order of magnitudes different from large-gene clusters. (C) and (D) display the fPC scores after filtering for very large genes (*>* 3000 bp) for replicates A and B.

**Figure A.3:**
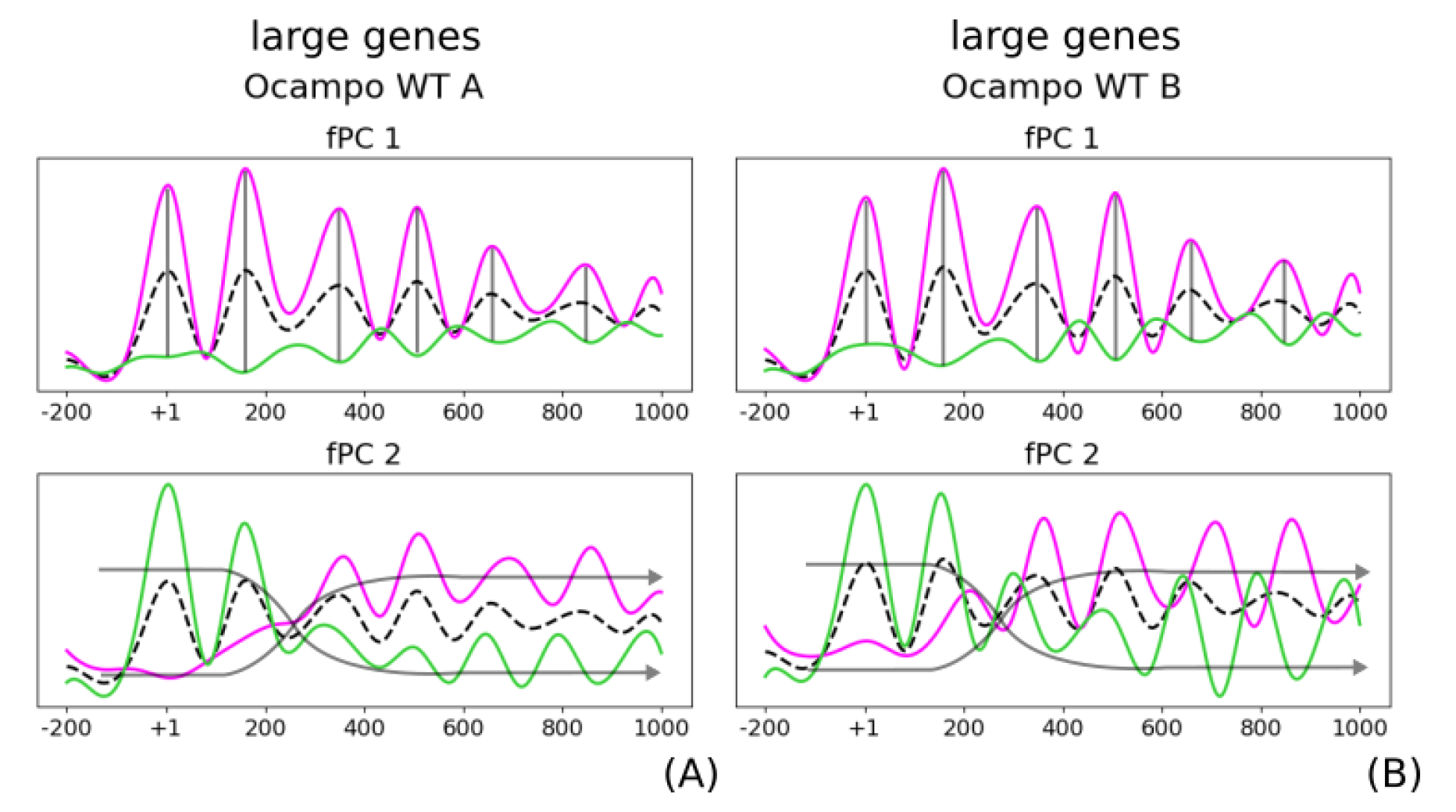
The large-gene fPC effect in WT. Despite fact that the functions differ in the A and B replicate ((A) and (B)), they both describe the same properties as when considering all genes (Fig 1(C)). To be precise, the first fPC describes seemingly position-dependent amplitude (dashed lines), and the second explains coordinated phasing (red arrows). The mean, a positive functional contribution, and a negative functional contribution are given in purple, green, and light blue, respectively.

**Figure A.4:**
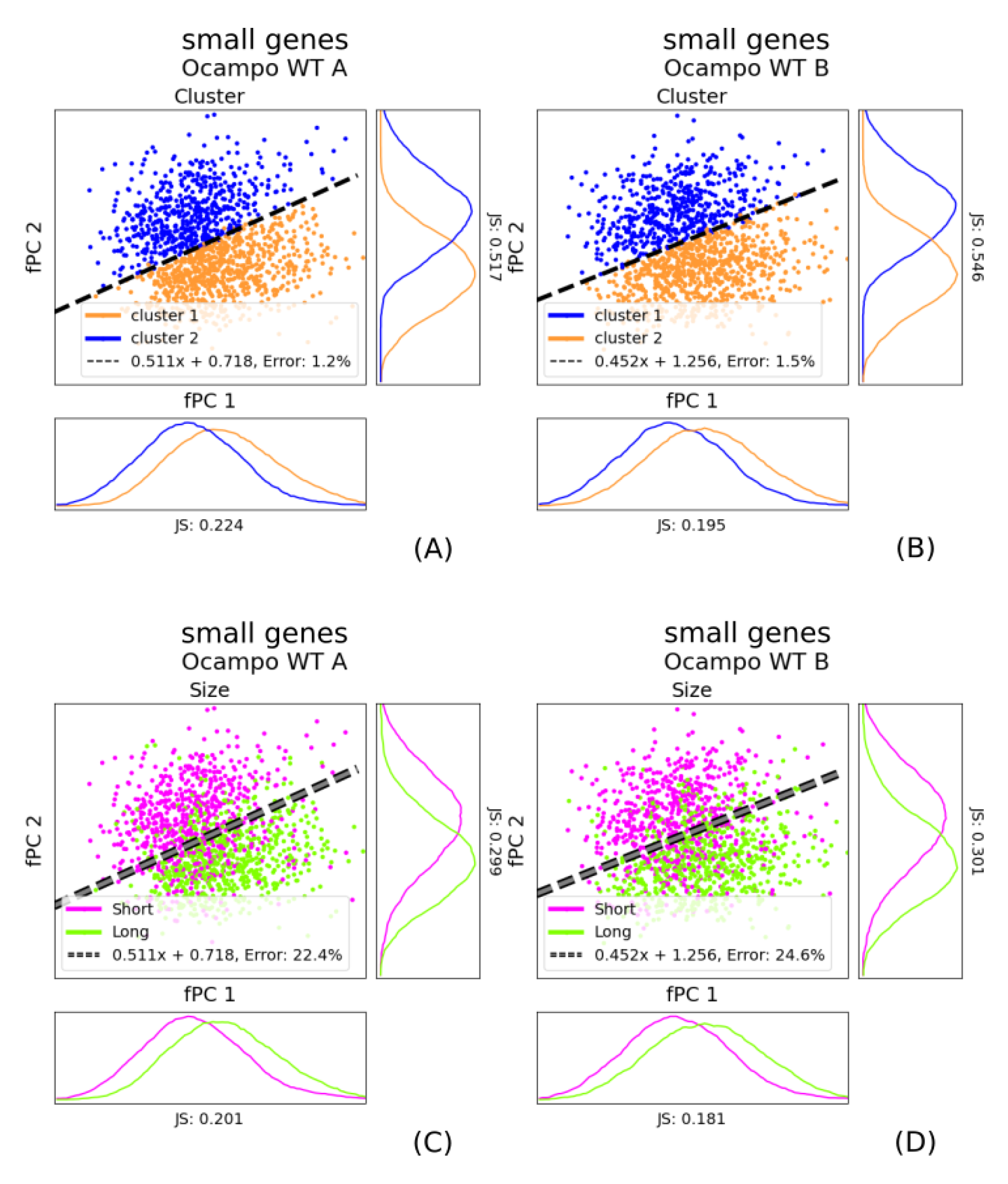
The Pearson coefficient clusters for exclusively small genes correspond to the gene size. When we repeated the Pearson coefficient clustering considering exclusively small genes, we can linearly separate again the two groups (orange and blue). However, this is predominantly explained by the size of the gene (short pink, long green). This in line with the hypothesis that coordinated nucleosome phasing along the transcribed region is strictly limited within the gene body. The phase separating line was determined on the Pearson clusters (dashed black line) using an SVM. The same separating boundary was also plotted in right plot showing grouping with respect to the size. We plotted the original SVM boundary from the Pearson clusters with a dashed grey line to indicate that it was not determined using gene size. (A) and (B) give the Pearson clusters for replicate A and B. (C) and (D) show the size dependence of replicate A and B.

**Figure A.5:**
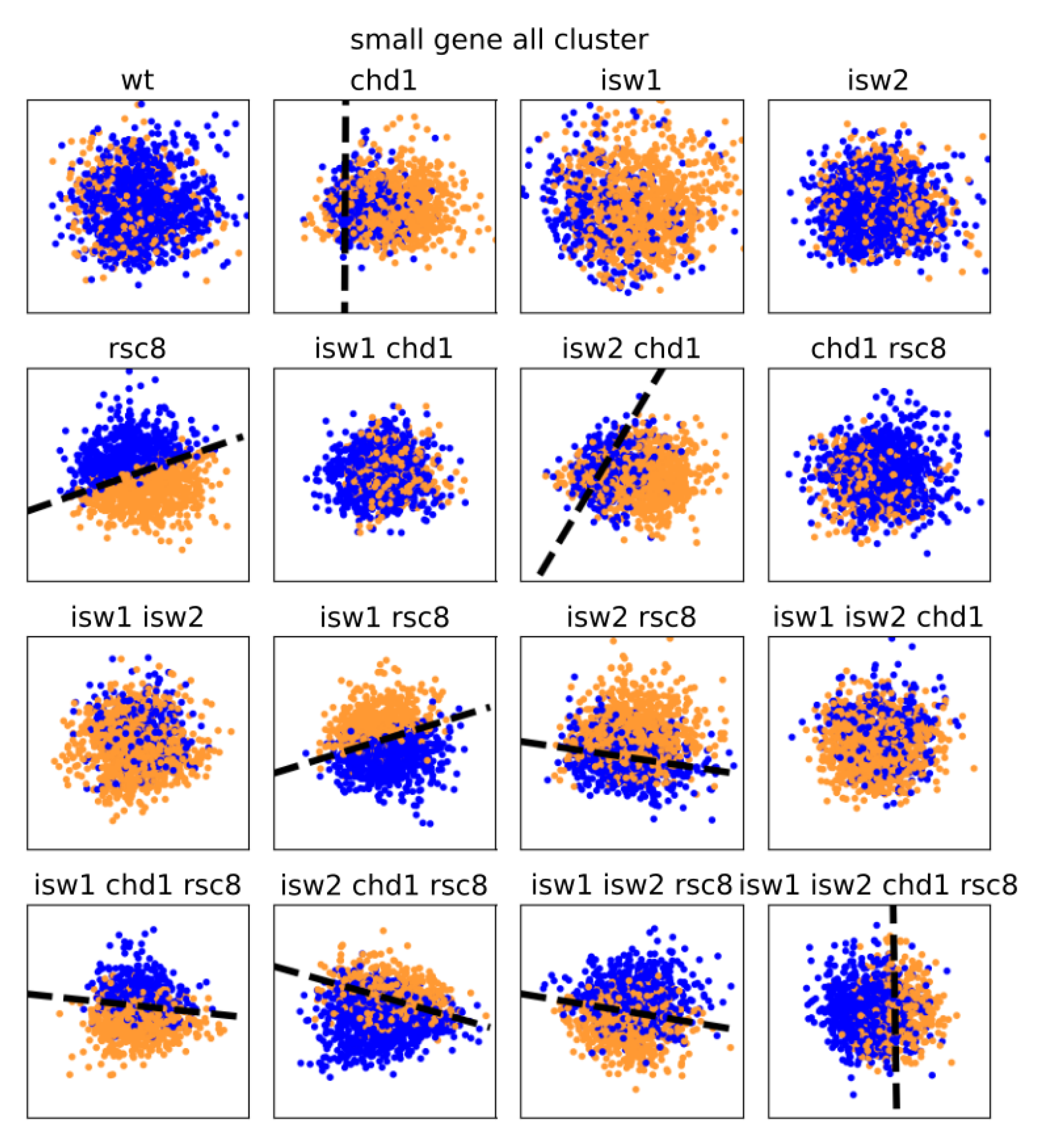
Pearson clusters of small genes lose separability with respect to their fPC scores. The figure shows the fPC scores *ζ* of small genes (*<* 1000 bp) of all conditions coloured with respect to the *all-gene* Pearson clustering. Green and red indicate each one group, the dashed line symbolises the best linear separation using a SVM. We removed the linear boundary in plots where it went through the periphery instead of dividing the data points. The x-axis represents the score of the first fPC *ζ*1, the y-axis gives the score for the second fPC *ζ*2. All axes are scaled to the same size; shapes are therefore comparable.

**Figure A.6:**
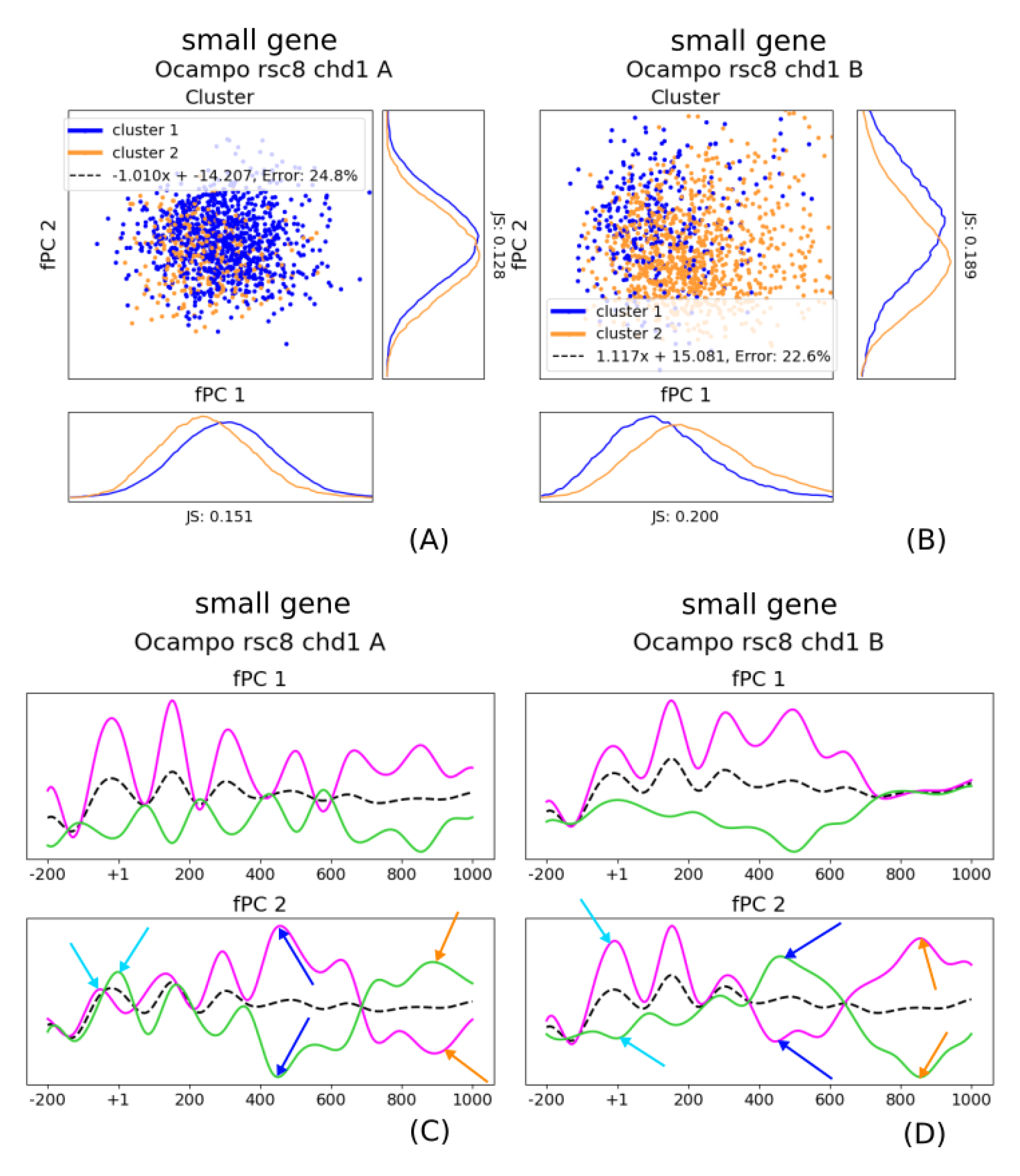
The small-gene fPC effect in *chd1*Δ*rsc8* strains. The double mutant seemingly re-establishes gene boundaries, and coordinated phasing is at least weakened after the +2 nucleosome (+1 in turquoise, +4 in blue, +6 in orange). This is true despite the fact that the A and B replicate differ. Figs (A) and (B) show the clusters for replicate *A* and *B*, and Figs (C) and (D) display their fPCs. We removed the separating boundaries in (A) and (B) because they did not reasonably divide the clusters. Nevertheless, we kept the estimated linear function in the legend to allow a comparison with other boundaries. Of particular note is the bias, which differs largely from large-gene clusters. the dashed black lines, the solid purple, and the solid green lines indicate the mean, a positive contribution, and a negative contribution, respectively.

**Figure A.7:**
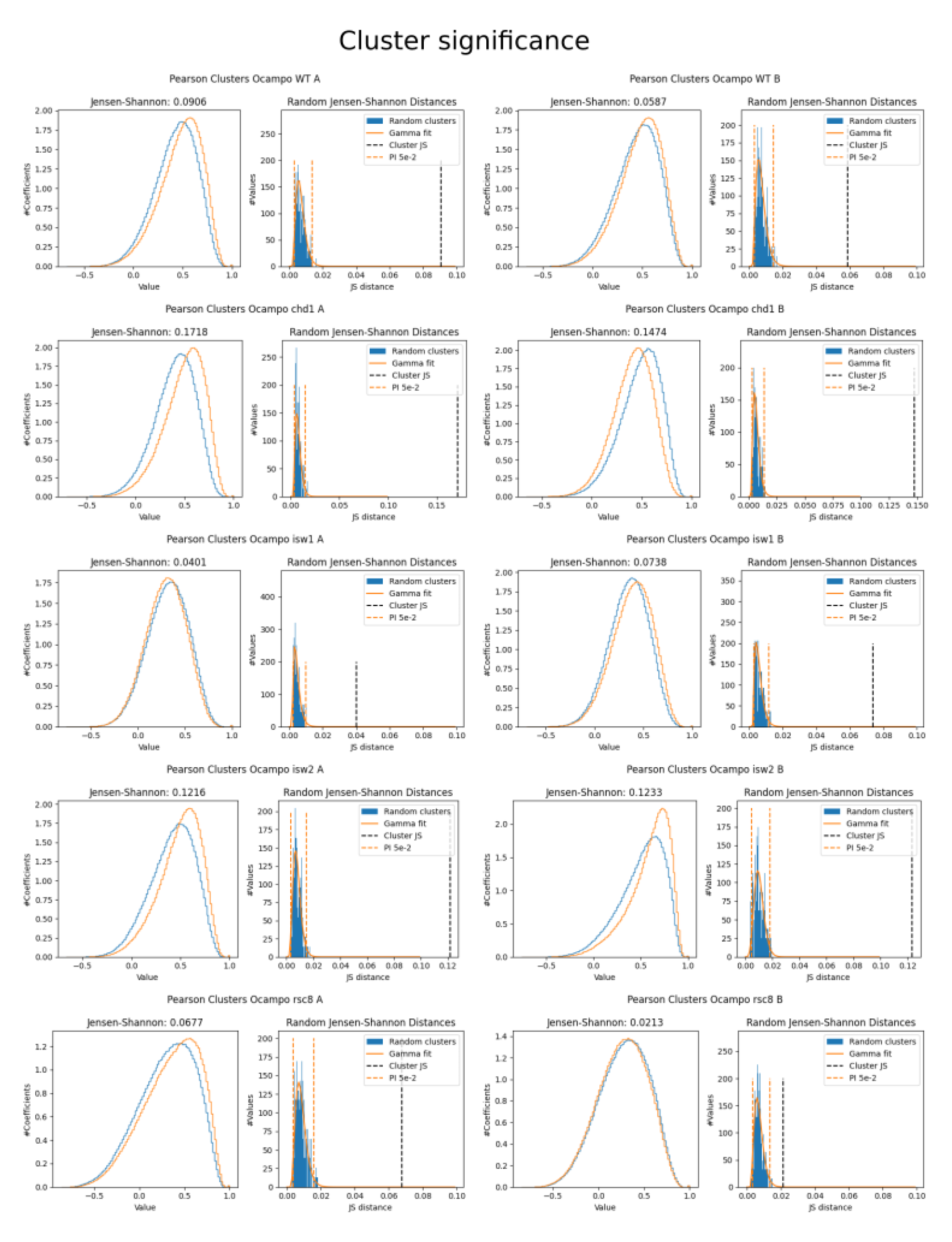
Cluster significance test for large genes (part 1). Left: The Pearson correlation coefficients for each profile in a cluster to all other distributions (independent of the cluster) is seemingly very similar for both groups, as indicated by the shape-independent the JS distance. Right: By measuring the JS for 500 random and mutually distinct clusters, we can approximate the expected distance over two random groupings of the Pearson coefficients using a gamma distribution. Orange dashed lines indicate the 95% PI, the dashed black line display the JS of the Pearson clusters.

**Figure A.7:**
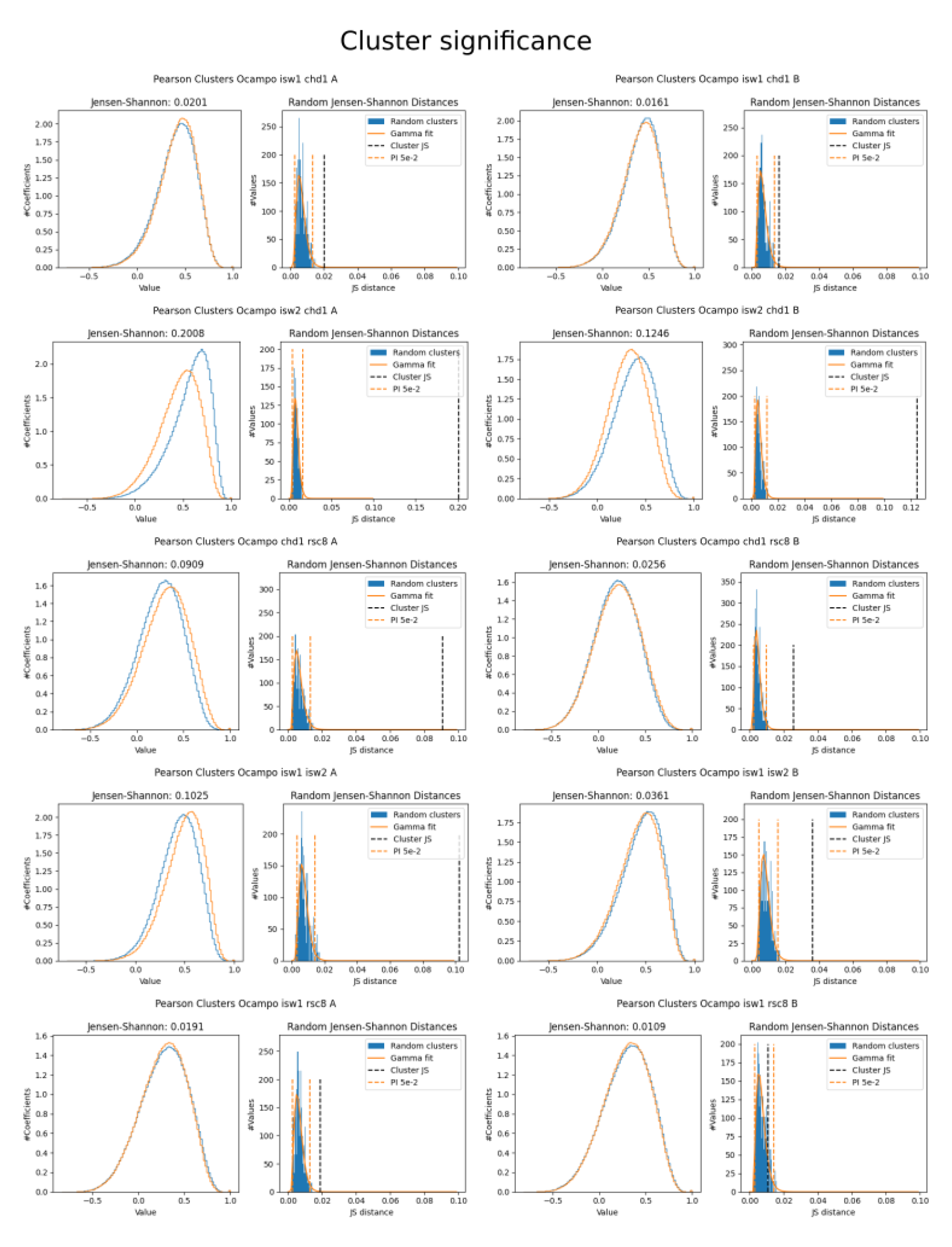
Cluster significance test for large genes (part 2). Left: The Pearson correlation coefficients for each profile in a cluster to all other distributions (independent of the cluster) is seemingly very similar for both groups, as indicated by the shape-independent the JS distance. Right: By measuring the JS for 500 random and mutually distinct clusters, we can approximate the expected distance over two random groupings of the Pearson coefficients using a gamma distribution. Orange dashed lines indicate the 95% PI, the dashed black line display the JS of the Pearson clusters.

**Figure A.7:**
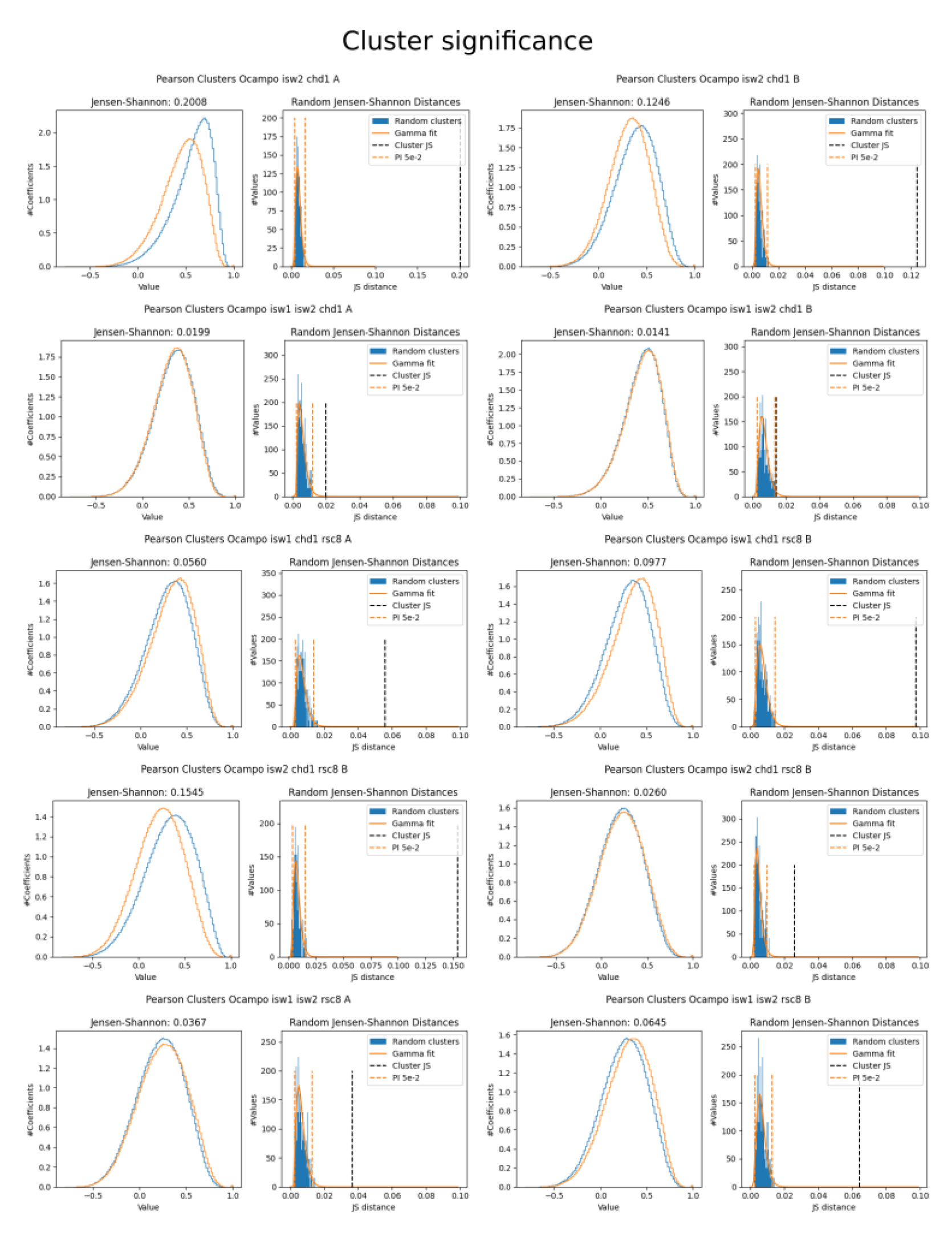
Cluster significance test for large genes (part 3). Left: The Pearson correlation coefficients for each profile in a cluster to all other distributions (independent of the cluster) is seemingly very similar for both groups, as indicated by the shape-independent the JS distance. Right: By measuring the JS for 500 random and mutually distinct clusters, we can approximate the expected distance over two random groupings of the Pearson coefficients using a gamma distribution. Orange dashed lines indicate the 95% PI, the dashed black line display the JS of the Pearson clusters.

**Figure A.7:**
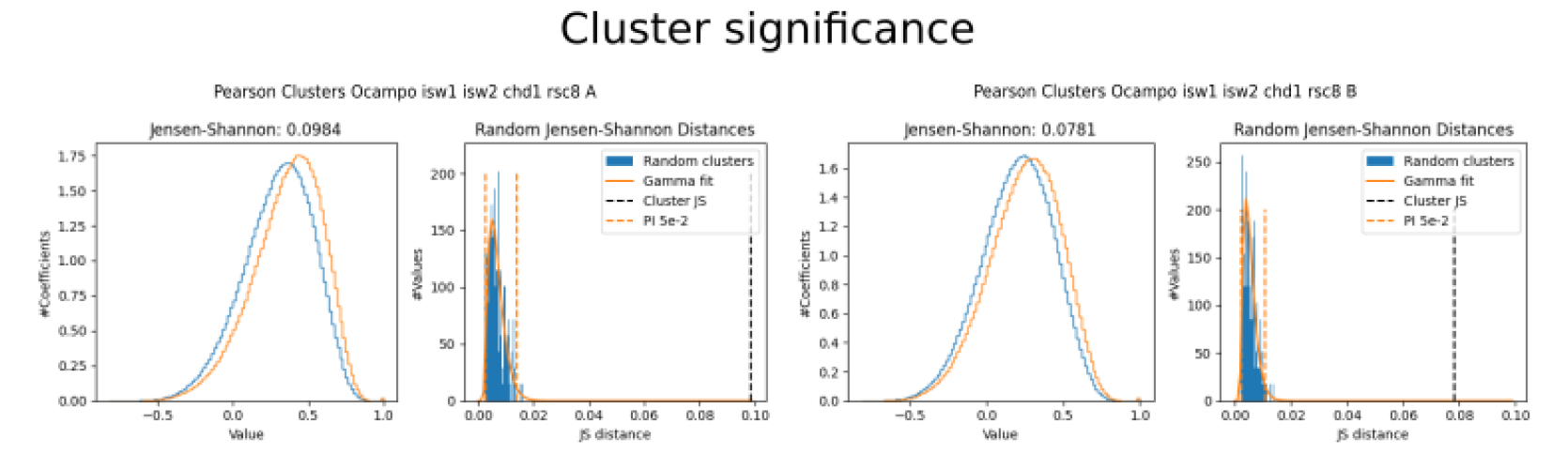
Cluster significance test for large genes (part 4). Left: The Pearson correlation coefficients for each profile in a cluster to all other distributions (independent of the cluster) is seemingly very similar for both groups, as indicated by the shape-independent the JS distance. Right: By measuring the JS for 500 random and mutually distinct clusters, we can approximate the expected distance over two random groupings of the Pearson coefficients using a gamma distribution. Orange dashed lines indicate the 95% PI, the dashed black line display the JS of the Pearson clusters.

**Figure A.8:**
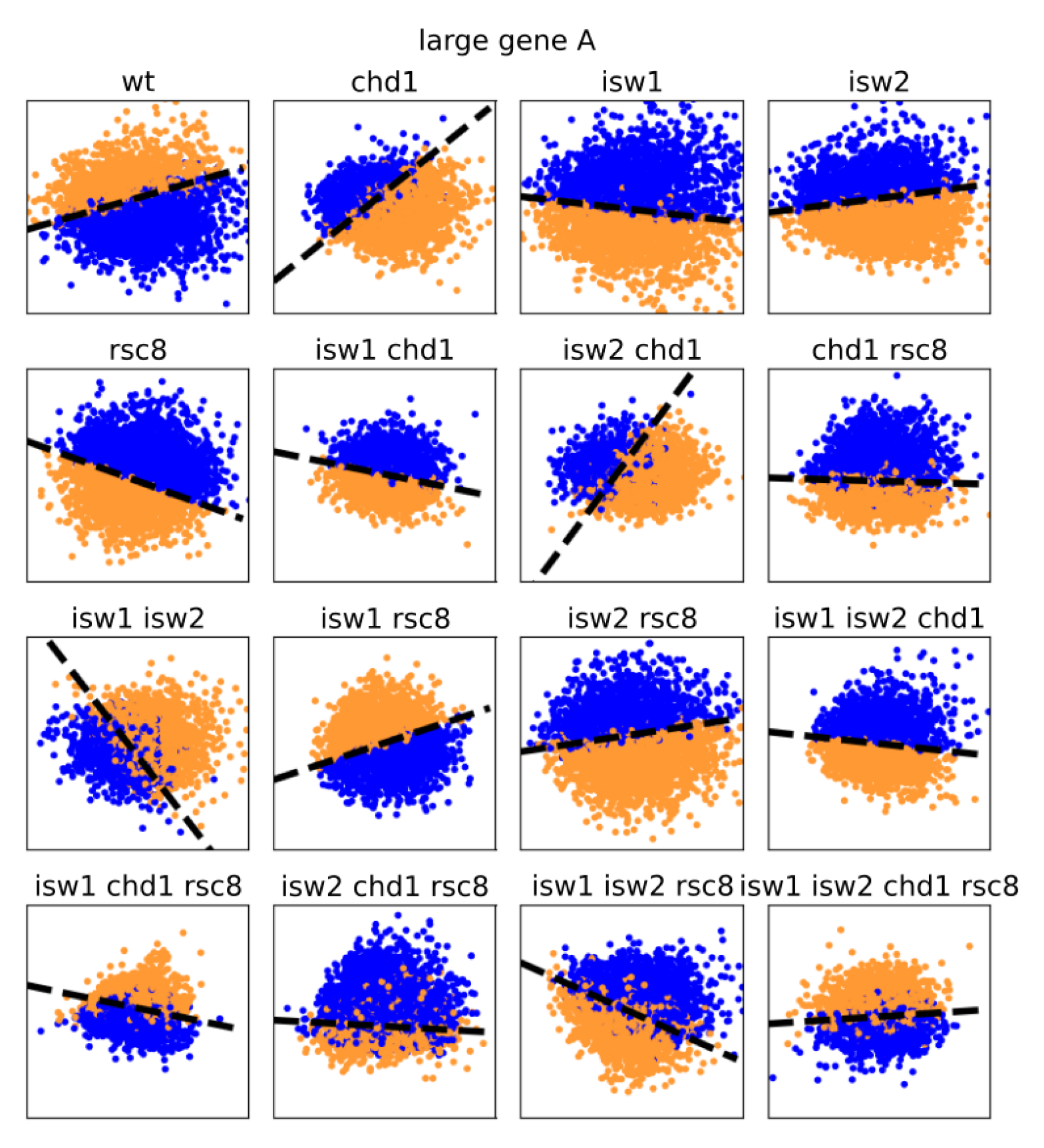
Pearson clusters of large genes are linearly separable with respect to their fPC scores (replicate A). The figure shows the fPC scores *ζ* of all conditions coloured with respect to the Pearson clustering using only large genes (*≥* 1000 bp). Blue and orange indicate each one group, the dashed line symbolises the best linear separation using a SVM. The x-axis represents the score of the first fPC *ζ*1, the y-axis gives the score for the second fPC *ζ*2. All axes are scaled to the same size; shapes are therefore comparable. It should be emphasised that only the absolute slope value matters and not the sign (i.e. pointing upwards or downwards).

**Figure A.9:**
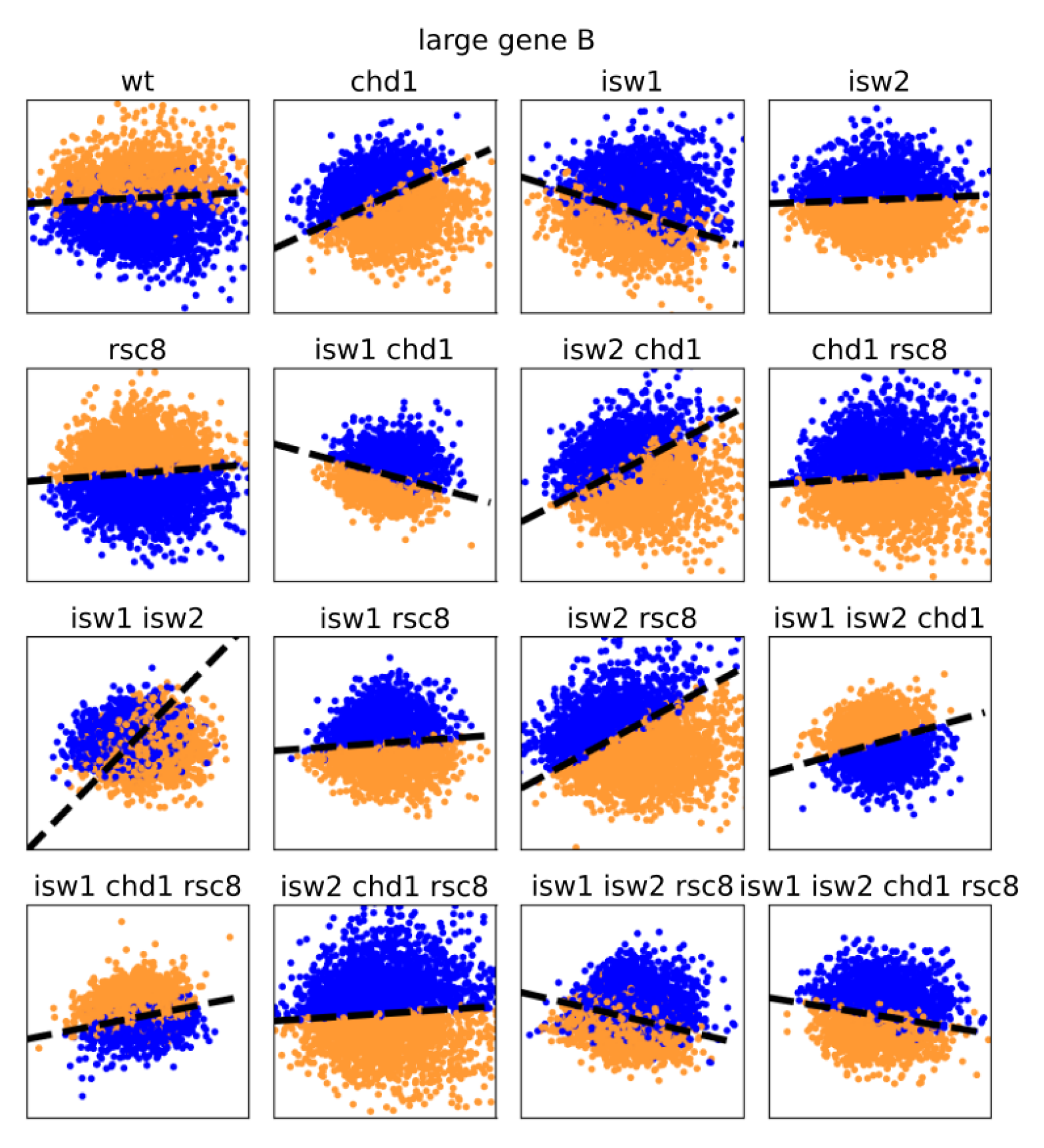
Pearson clusters of large genes are linearly separable with respect to their fPC scores (replicate B). The figure shows the fPC scores *ζ* of all conditions coloured with respect to the Pearson clustering using only large genes (*≥* 1000 bp). Blue and orange indicate each one group, the dashed line symbolises the best linear separation using a SVM. The x-axis represents the score of the first fPC *ζ*1, the y-axis gives the score for the second fPC *ζ*2. All axes are scaled to the same size; shapes are therefore comparable. It should be emphasised that only the absolute slope value matters and not the sign (i.e. pointing upwards or downwards).

**Figure A.10:**
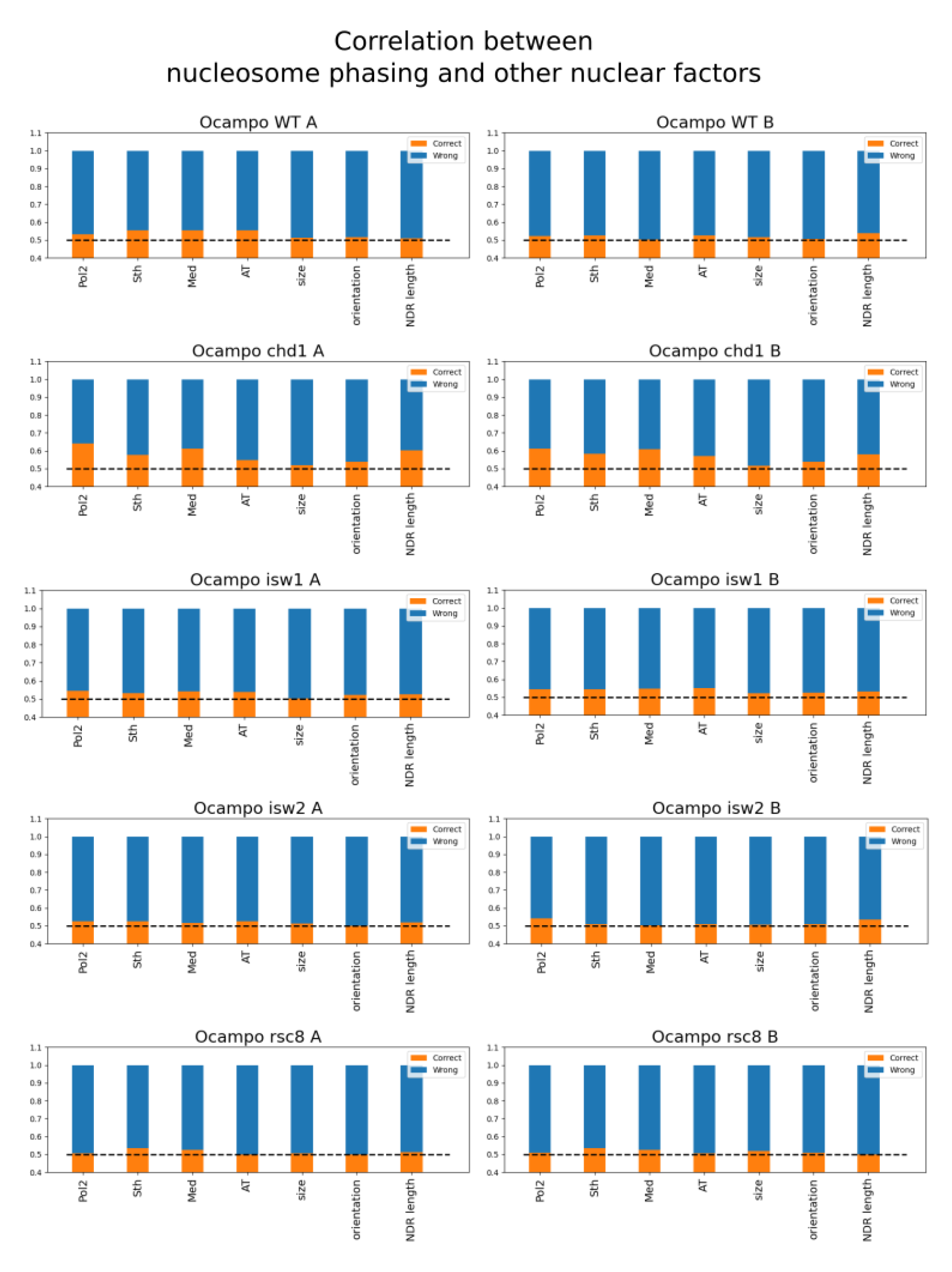
Interdependence of Pearson clusters of MNase-seq profiles and other nuclear factors (part 1). The orange bar shows the ratio of cases where the nuclear factor could predict clustering, blue gives the wrongly classified ratio. Random guessing would be correct in 50% of the cases, which is given by the dashed black line. Consequently, the orange bar must exceed the dashed line to suggest interdependence.

**Figure A.10:**
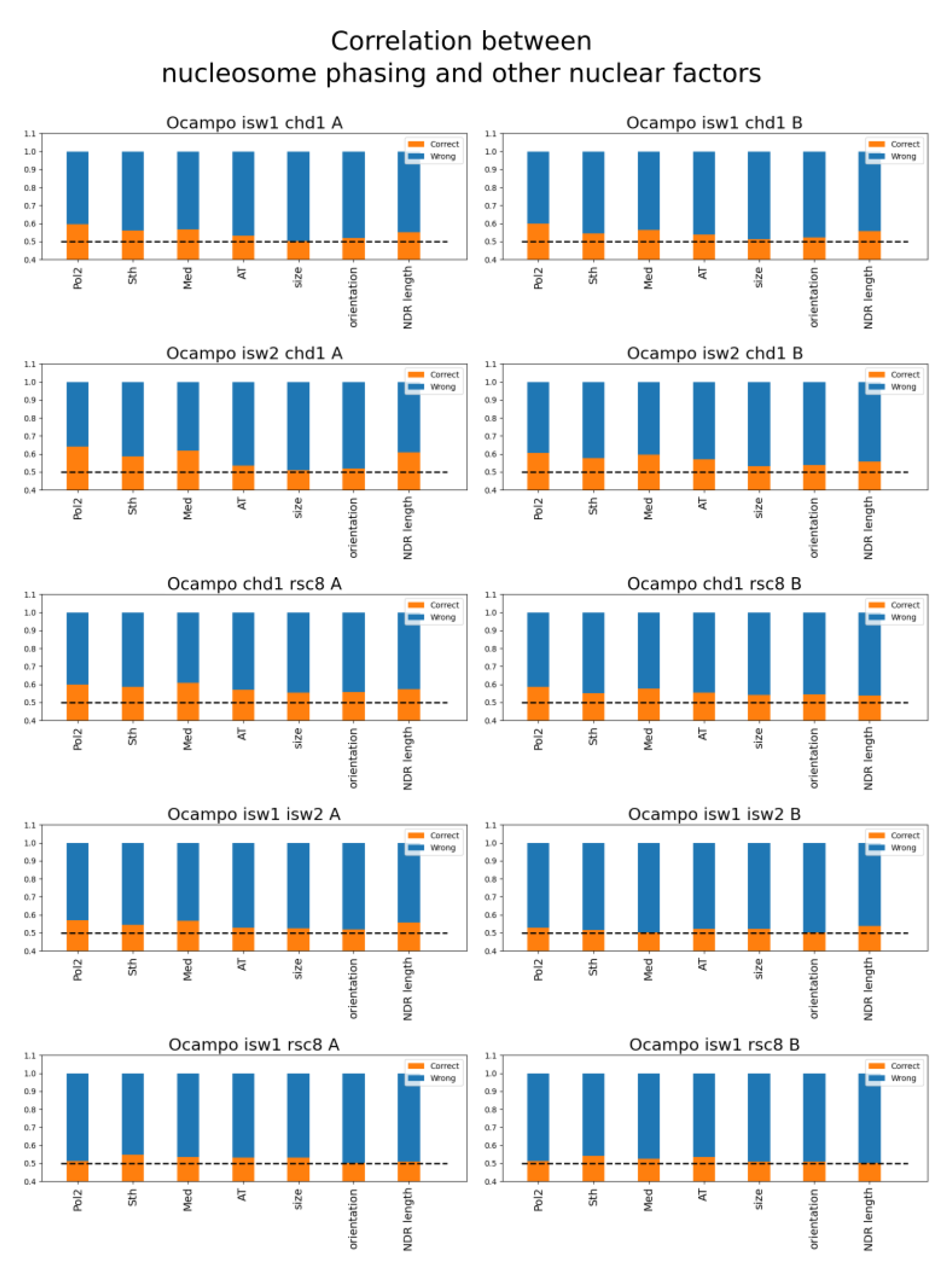
Interdependence of Pearson clusters of MNase-seq profiles and other nuclear factors (part 2). The orange bar shows the ratio of cases where the nuclear factor could predict clustering, blue gives the wrongly classified ratio. Random guessing would be correct in 50% of the cases, which is given by the dashed black line. Consequently, the orange bar must exceed the dashed line to suggest interdependence.

**Figure A.10:**
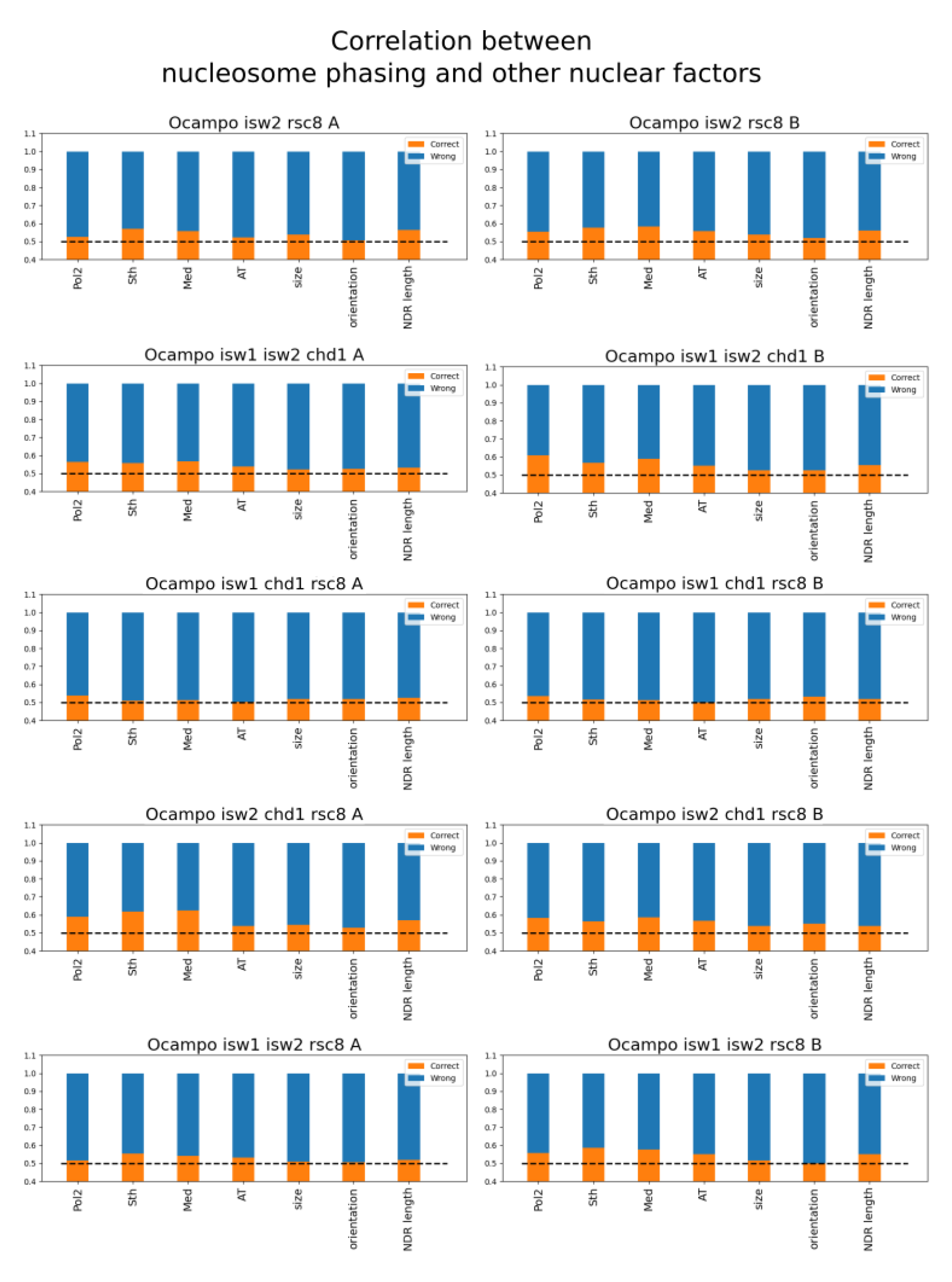
Interdependence of Pearson clusters of MNase-seq profiles and other nuclear factors (part 3). The orange bar shows the ratio of cases where the nuclear factor could predict clustering, blue gives the wrongly classified ratio. Random guessing would be correct in 50% of the cases, which is given by the dashed black line. Consequently, the orange bar must exceed the dashed line to suggest interdependence.

**Figure A.10:**
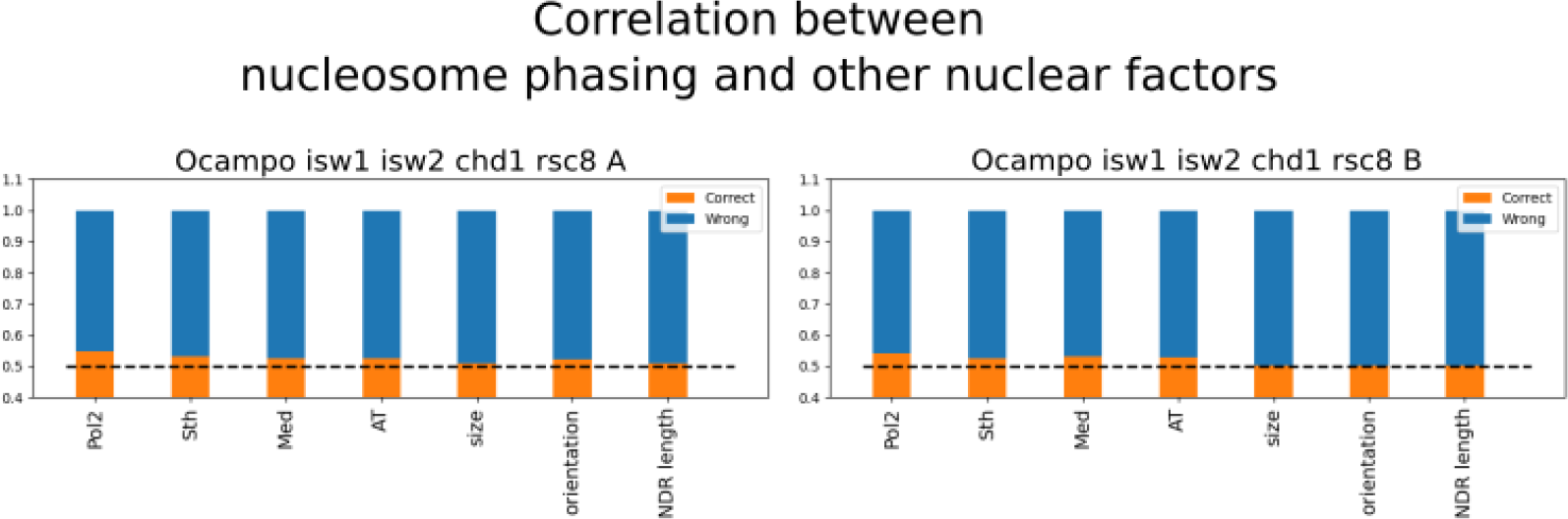
Interdependence of Pearson clusters of MNase-seq profiles and other nuclear factors (part 4). The orange bar shows the ratio of cases where the nuclear factor could predict clustering, blue gives the wrongly classified ratio. Random guessing would be correct in 50% of the cases, which is given by the dashed black line. Consequently, the orange bar must exceed the dashed line to suggest interdependence.

**Figure A.11:**
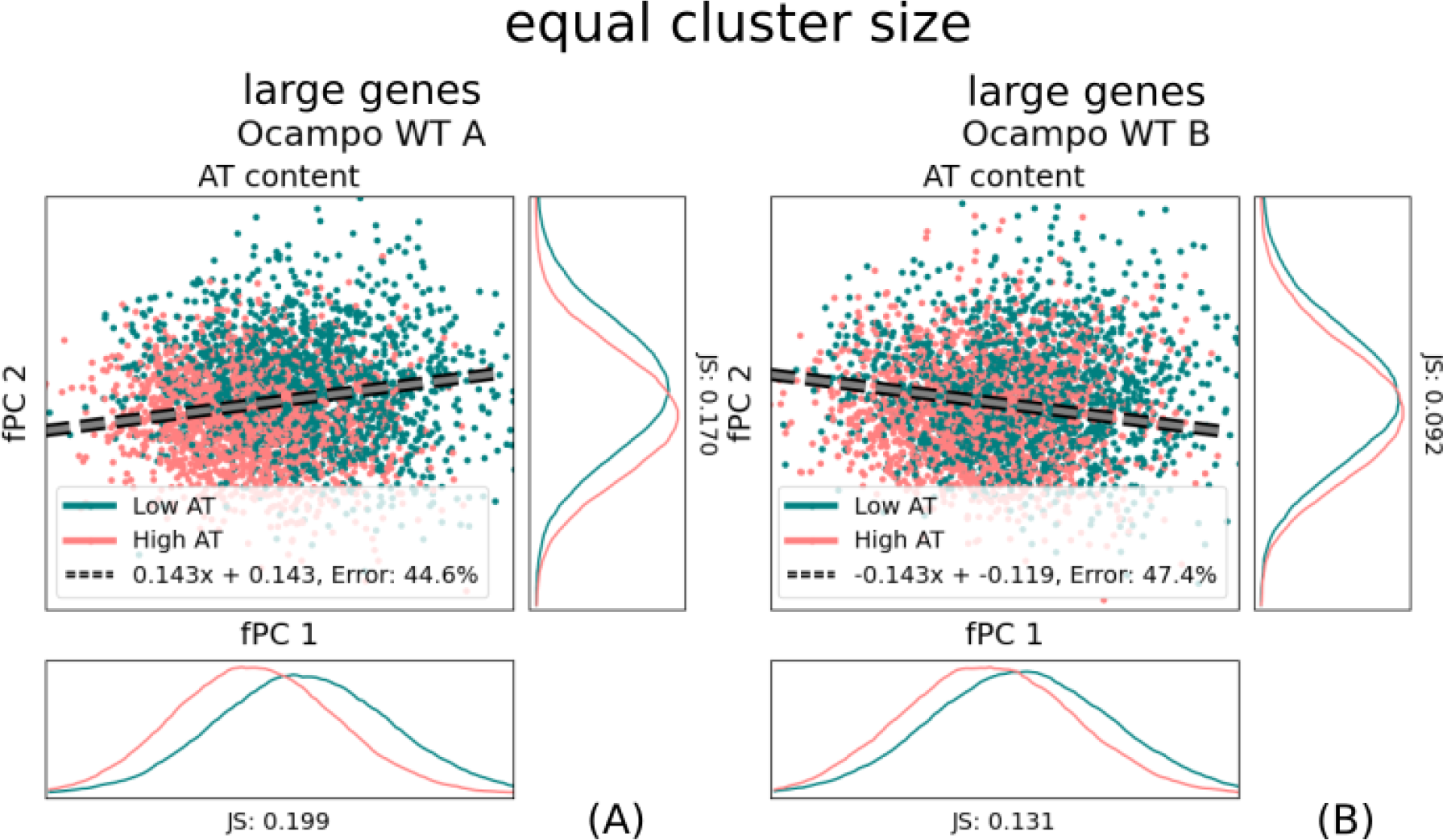
AT-ratio distribution with respect to the fPC scores. Whilst there is seemingly a slight correlation between Pearson coefficient clusters and AT-ratio in the *A* replicate, this is trend vanishes for the *B* replicate. In fact, both replicates might rather distribute AT-rich and AT-poor genes orthogonal to the dividing boundary. We plotted the original SVM boundary from the Pearson clusters with a dashed grey line to indicate that it was not determined using the AT content.

## References

1. Roger Kornberg. The location of nucleosomes in chromatin: specific or statistical? Nature, 292:579–580, 1981.

2. Travis N Mavrich, Ilya P Ioshikhes, Bryan J Venters, Cizhong Jiang, Lynn P Tomsho, Ji Qi, Stephan C Schuster, Istvan Albert, and B Franklin Pugh. A barrier nucleosome model for statistical positioning of nucleosomes throughout the yeast genome. Genome research, 18(7):1073–1083, 2008.

3. Kevin Struhl and Eran Segal. Determinants of nucleosome positioning. Nature structural & molecular biology, 20(3):267–273, 2013.

4. Yong Zhang, Zarmik Moqtaderi, Barbara P Rattner, Ghia Euskirchen, Michael Snyder, James T Kadonaga, X Shirley Liu, and Kevin Struhl. Intrinsic histone-dna interactions are not the major determinant of nucleosome positions in vivo. Nature structural & molecular biology, 16(8):847–852, 2009.

5. Pauline Vasseur, Saphia Tonazzini, Rahima Ziane, Alain Camasses, Oliver J Rando, and Marta Radman-Livaja. Dynamics of nucleosome positioning maturation following genomic replication. Cell reports, 16(10):2651–2665, 2016.

6. Amanda L Hughes, Yi Jin, Oliver J Rando, and Kevin Struhl. A functional evolutionary approach to identify determinants of nucleosome positioning: a unifying model for establishing the genome-wide pattern. Molecular cell, 48(1):5–15, 2012.

7. Job Dekker. Gc-and at-rich chromatin domains differ in conformation and histone modification status and are differentially modulated by rpd3p. Genome biology, 8:1–14, 2007.

8. Alberto Marin-Gonzalez, JG Vilhena, Ruben Perez, and Fernando Moreno-Herrero. A molecular view of dna flexibility. Quarterly Reviews of Biophysics, 54:e8, 2021.

9. Cedric R Clapier, Janet Iwasa, Bradley R Cairns, and Craig L Peterson. Mechanisms of action and regulation of atp-dependent chromatin-remodelling complexes. Nature reviews Molecular cell biology, 18(7):407–422, 2017.

10. Timothy J Parnell, Jason T Huff, and Bradley R Cairns. Rsc regulates nucleosome positioning at pol ii genes and density at pol iii genes. The EMBO journal, 27(1):100–110, 2008.

11. Carlo Yague-Sanz, Enrique Vázquez, Mar Sánchez, Francisco Antequera, and Damien Hermand. A conserved role of the rsc chromatin remodeler in the establishment of nucleosome-depleted regions. Current Genetics, 63:187–193, 2017.

12. Josefina Ocampo, Răzvan V Chereji, Peter R Eriksson, and David J Clark. Contrasting roles of the rsc and isw1/chd1 chromatin remodelers in rna polymerase ii elongation and termination. Genome research, 29(3):407–417, 2019.

13. Emily Biernat, Jeena Kinney, Kyle Dunlap, Christian Rizza, and Chhabi K Govind. The rsc complex remodels nucleosomes in transcribed coding sequences and promotes transcription in saccharomyces cerevisiae. Genetics, 217(4):iyab021, 2021.

14. Dwaipayan Ganguli, Razvan V Chereji, James R Iben, Hope A Cole, and David J Clark. Rsc-dependent constructive and destructive interference between opposing arrays of phased nucleosomes in yeast. Genome research, 24(10):1637–1649, 2014.

15. Manu Shubhdarshan Shukla, Sajad Hussain Syed, Fabien Montel, Cendrine Faivre-Moskalenko, Jan Bednar, Andrew Travers, Dimitar Angelov, and Stefan Dimitrov. Remosomes: Rsc generated non-mobilized particles with approximately 180 bp dna loosely associated with the histone octamer. Proceedings of the National Academy of Sciences, 107(5):1936–1941, 2010.

16. Claudia Alén, Nicholas A Kent, Hannah S Jones, Justin O’Sullivan, Agustın Aranda, and Nicholas J Proudfoot. A role for chromatin remodeling in transcriptional termination by rna polymerase ii. Molecular cell, 10(6):1441–1452, 2002.

17. Marta Radman-Livaja, Tiffani K Quan, Lourdes Valenzuela, Jennifer A Armstrong, Tibor Van Welsem, TaeSoo Kim, Laura J Lee, Stephen Buratowski, Fred Van Leeuwen, Oliver J Rando, et al. A key role for chd1 in histone h3 dynamics at the 3’ ends of long genes in yeast. PLoS genetics, 8(7):e1002811, 2012.

18. Josefina Ocampo, Răzvan V Chereji, Peter R Eriksson, and David J Clark. The isw1 and chd1 atp-dependent chromatin remodelers compete to set nucleosome spacing in vivo. Nucleic acids research, 44(10):4625–4635, 2016.

19. Triantaffyllos Gkikopoulos, Pieta Schofield, Vijender Singh, Marina Pinskaya, Jane Mellor, Michaela Smolle, Jerry L Workman, Geoffrey J Barton, and Tom Owen-Hughes. A role for snf2-related nucleosome-spacing enzymes in genome-wide nucleosome organization. Science, 333(6050):1758–1760, 2011.

20. Toshio Tsukiyama, Jeffrey Palmer, Carolyn C Landel, Joseph Shiloach, and Carl Wu. Characterization of the imitation switch subfamily of atp-dependent chromatin-remodeling factors in saccharomyces cerevisiae. Genes & development, 13(6):686–697, 1999.

21. Iestyn Whitehouse, Oliver J Rando, Jeff Delrow, and Toshio Tsukiyama. Chromatin remodelling at promoters suppresses antisense transcription. Nature, 450(7172):1031–1035, 2007.

22. Janet G Yang, Tina Shahian Madrid, Elena Sevastopoulos, and Geeta J Narlikar. The chromatin-remodeling enzyme acf is an atp-dependent dna length sensor that regulates nucleosome spacing. Nature structural & molecular biology, 13(12):1078–1083, 2006.

23. Kazuhiro Yamada, Timothy D Frouws, Brigitte Angst, Daniel J Fitzgerald, Carl DeLuca, Kyoko Schimmele, David F Sargent, and Timothy J Richmond. Structure and mechanism of the chromatin remodelling factor isw1a. Nature, 472(7344):448–453, 2011.

24. Corinna Lieleg, Philip Ketterer, Johannes Nuebler, Johanna Ludwigsen, Ulrich Gerland, Hendrik Dietz, Felix Mueller-Planitz, and Philipp Korber. Nucleosome spacing generated by iswi and chd1 remodelers is constant regardless of nucleosome density. Molecular and cellular biology, 35(9):1588–1605, 2015.

25. Elisa Oberbeckmann, Vanessa Niebauer, Shinya Watanabe, Lucas Farnung, Manuela Moldt, Andrea Schmid, Patrick Cramer, Craig L Peterson, Sebastian Eustermann, Karl-Peter Hopfner, et al. Ruler elements in chromatin remodelers set nucleosome array spacing and phasing. Nature Communications, 12(1):3232, 2021.

26. Cizhong Jiang and B Franklin Pugh. Nucleosome positioning and gene regulation: advances through genomics. Nature Reviews Genetics, 10(3):161–172, 2009.

27. Assaf Weiner, Amanda Hughes, Moran Yassour, Oliver J Rando, and Nir Friedman. High-resolution nucleosome mapping reveals transcription-dependent promoter packaging. Genome research, 20(1):90–100, 2010.

28. R Thomas Koerber, Ho Sung Rhee, Cizhong Jiang, and B Franklin Pugh. Interaction of transcriptional regulators with specific nucleosomes across the saccharomyces genome. Molecular cell, 35(6):889–902, 2009.

29. Yunhao Wang, Anqi Wang, Zujun Liu, Andrew L Thurman, Linda S Powers, Meng Zou, Yue Zhao, Adam Hefel, Yunyi Li, Joseph Zabner, et al. Single-molecule long-read sequencing reveals the chromatin basis of gene expression. Genome research, 29(8):1329–1342, 2019.

30. Ashish Kumar Singh, Tamás Schauer, Lena Pfaller, Tobias Straub, and Felix Mueller-Planitz. The biogenesis and function of nucleosome arrays. Nature communications, 12(1):7011, 2021.

31. Sandro Baldi, Stefan Krebs, Helmut Blum, and Peter B Becker. Genome-wide measurement of local nucleosome array regularity and spacing by nanopore sequencing. Nature structural & molecular biology, 25(9):894–901, 2018.

32. Binbin Lai, Weiwu Gao, Kairong Cui, Wanli Xie, Qingsong Tang, Wenfei Jin, Gangqing Hu, Bing Ni, and Keji Zhao. Principles of nucleosome organization revealed by single-cell micrococcal nuclease sequencing. Nature, 562(7726):281–285, 2018.

33. Jun Wan, Jimmy Lin, Donald J Zack, and Jiang Qian. Relating periodicity of nucleosome organization and gene regulation. Bioinformatics, 25(14):1782–1788, 2009.

34. Răzvan V Chereji and David J Clark. Major determinants of nucleosome positioning. Biophysical journal, 114(10):2279–2289, 2018.

35. Yanghui Liu, Yehua Li, Raymond J Carroll, and Naisyin Wang. Predictive functional linear models with diverging number of semiparametric single-index interactions. Journal of Econometrics, 230(2):221–239, 2022.

36. Reka Karuppusami, Belavendra Antonisamy, and Prasanna S Premkumar. Functional principal component analysis for identifying the child growth pattern using longitudinal birth cohort data. BMC Medical Research Methodology, 22(1):76, 2022.

37. Li Luo, Yun Zhu, and Momiao Xiong. Smoothed functional principal component analysis for testing association of the entire allelic spectrum of genetic variation. European Journal of Human Genetics, 21(2):217–224, 2013.

38. Daechan Park, Adam R Morris, Anna Battenhouse, and Vishwanath R Iyer. Simultaneous mapping of transcript ends at single-nucleotide resolution and identification of widespread promoter-associated non-coding rna governed by tata elements. Nucleic acids research, 42(6):3736–3749, 2014.

39. Avital Klein-Brill, Daphna Joseph-Strauss, Alon Appleboim, and Nir Friedman. Dynamics of chromatin and transcription during transient depletion of the rsc chromatin remodeling complex. Cell reports, 26(1):279–292, 2019.

40. Donald O Hebb. Organization of behavior. *New York*: Wiley and Sons. J. Clin. Psychology, 6(3):335–307, 1949.

41. Slawomir Kubik, Maria Jessica Bruzzone, Philippe Jacquet, Jean-Luc Falcone, Jacques Rougemont, and David Shore. Nucleosome stability distinguishes two different promoter types at all protein-coding genes in yeast. Molecular cell, 60(3):422–434, 2015.

42. Bradley R Cairns, Yahli Lorch, Yang Li, Mincheng Zhang, Lynne Lacomis, Hediye Erdjument-Bromage, Paul Tempst, Jian Du, Brehon Laurent, and Roger D Kornberg. Rsc, an essential, abundant chromatin-remodeling complex. Cell, 87(7):1249–1260, 1996.

43. Nils Krietenstein, Megha Wal, Shinya Watanabe, Bongsoo Park, Craig L Peterson, B Franklin Pugh, and Philipp Korber. Genomic nucleosome organization reconstituted with pure proteins. Cell, 167(3):709–721, 2016.

44. Marc Damelin, Itamar Simon, Terence I Moy, Boris Wilson, Suzanne Komili, Paul Tempst, Frederick P Roth, Richard A Young, Bradley R Cairns, and Pamela A Silver. The genome-wide localization of rsc9, a component of the rsc chromatin-remodeling complex, changes in response to stress. Molecular cell, 9(3):563–573, 2002.

45. Huck Hui Ng, Fraņcois Robert, Richard A Young, and Kevin Struhl. Genome-wide location and regulated recruitment of the rsc nucleosome-remodeling complex. Genes & development, 16(7):806–819, 2002.

46. Tiffani Kiyoko Quan and Grant Ashley Hartzog. Histone h3k4 and k36 methylation, chd1 and rpd3s oppose the functions of saccharomyces cerevisiae spt4–spt5 in transcription. Genetics, 184(2):321–334, 2010.

47. Daechan Park, Haridha Shivram, and Vishwanath R Iyer. Chd1 co-localizes with early transcription elongation factors independently of h3k36 methylation and releases stalled rna polymerase ii at introns. Epigenetics & chromatin, 7(1):1–11, 2014.

48. Rajna Simic, Derek L Lindstrom, Hien G Tran, Kelli L Roinick, Patrick J Costa, Alexander D Johnson, Grant A Hartzog, and Karen M Arndt. Chromatin remodeling protein chd1 interacts with transcription elongation factors and localizes to transcribed genes. The EMBO journal, 22(8):1846–1856, 2003.

49. Concetta GA Marfella and Anthony N Imbalzano. The chd family of chromatin remodelers. Mutation Research/Fundamental and Molecular Mechanisms of Mutagenesis, 618(1-2):30–40, 2007.

50. Kévin M André, Nathalie Giordanengo Aiach, Veronica Martinez-Fernandez, Leo Zeitler, Adriana Alberti, Arach Goldar, Michel Werner, Cyril Denby Wilkes, and Julie Soutourina. Functional interplay between mediator and rsc chromatin remodeling complex controls nucleosome-depleted region maintenance at promoters. Cell Reports, 42(5), 2023.

51. Felix Krueger. Trim galore. https://github.com/FelixKrueger/TrimGalore/releases/tag/0.6.5, 2019.

52. Marcel Martin. Cutadapt removes adapter sequences from high-throughput sequencing reads. *EMBnet*. journal, 17(1):10–12, 2011.

53. Ben Langmead and Steven L Salzberg. Fast gapped-read alignment with bowtie 2. Nature methods, 9(4):357–359, 2012.

54. Heng Li, Bob Handsaker, Alec Wysoker, Tim Fennell, Jue Ruan, Nils Homer, Gabor Marth, Goncalo Abecasis, Richard Durbin, and 1000 Genome Project Data Processing Subgroup. The sequence alignment/map format and samtools. bioinformatics, 25(16):2078–2079, 2009.

55. Fidel Ramírez, Devon P Ryan, Björn Gruüing, Vivek Bhardwaj, Fabian Kilpert, Andreas S Richter, Steffen Heyne, Friederike Duüdar, and Thomas Manke. deeptools2: a next generation web server for deep-sequencing data analysis. Nucleic acids research, 44(W1):W160–W165, 2016.

56. Adrien Georges, Diyavarshini Gopaul, Cyril Denby Wilkes, Nathalie Giordanengo Aiach, Elizaveta Novikova, Marie-Bénédicte Barrault, Olivier Alibert, and Julie Soutourina. Functional interplay between mediator and rna polymerase ii in rad2/xpg loading to the chromatin. Nucleic acids research, 47(17):8988–9004, 2019.

57. Leonard Kaufman and Peter J Rousseeuw. Finding groups in data: an introduction to cluster analysis. John Wiley & Sons, 2009.

58. Peter J Rousseeuw. Silhouettes: a graphical aid to the interpretation and validation of cluster analysis. Journal of computational and applied mathematics, 20:53–65, 1987.

59. Carlos Ramos-Carreño, José Luis Torrecilla, Miguel Carbajo-Berrocal, Pablo Marcos, and Alberto Suárez. scikit-fda: a python package for functional data analysis. arXiv preprint arXiv*:2211.02566*, 2022.

